# Probing the Hidden Sensitivity of Intrinsically Disordered Proteins to their Chemical Environment

**DOI:** 10.1101/2020.08.17.252478

**Authors:** David Moses, Feng Yu, Garrett M. Ginell, Nora M. Shamoon, Patrick S. Koenig, Alex S. Holehouse, Shahar Sukenik

## Abstract

Intrinsically disordered proteins and protein-regions (IDRs) make up roughly 30% of the human proteome and play vital roles in a wide variety of biological processes. Given a lack of persistent tertiary structure, all of the residues in an IDR are, to some extent, solvent exposed. This extensive surface area, coupled with the absence of strong intramolecular contacts, makes IDRs inherently sensitive to their chemical environment. Despite this sensitivity, our understanding of how IDR structural ensembles are influenced by changes in their chemical environment is limited. This is particularly relevant given a growing body of evidence showing that IDR function is linked to the underlying structural ensemble. We develop and use a combined experimental, computational, and analytical framework for high-throughput characterization of IDR sensitivity we call solution space scanning. Our framework reveals that IDRs show sequence-dependent sensitivity to solution chemistry, with complex behavior that can be interpreted through relatively simple polymer models. Our results imply that solution-responsive IDRs are ubiquitous and can provide an additional layer of biological regulation.

---

Intrinsically disordered proteins and protein-regions (IDRs) play key roles in mediating cellular signalling, transcriptional regulation, and homeostatic functions^1^. IDRs differ from well-folded proteins in that they exist in an ensemble of rapidly changing configurations (**Fig. 1a**). This conformational ensemble is often tied to IDR function^2–4^. IDR ensembles have extensive surface area exposed to the surrounding solution, and few non-covalent intramolecular bonds that constrain their structure. As such, IDR ensembles are highly malleable and can be strongly affected by the chemistry of their surrounding environment^5^. Inside the cell, the chemical composition can change due to routine cell-cycle events or external stress^6–10^. The plasticity of IDR ensembles makes them ideal sensors and actuators of these changes^11–13^, but perhaps also impairs their activity in deleterious environments such as metabolically rewired cancer cells^14^. Still, little effort has been directed at systematically characterizing IDR sensitivity to solution changes.

**Figure 1.**
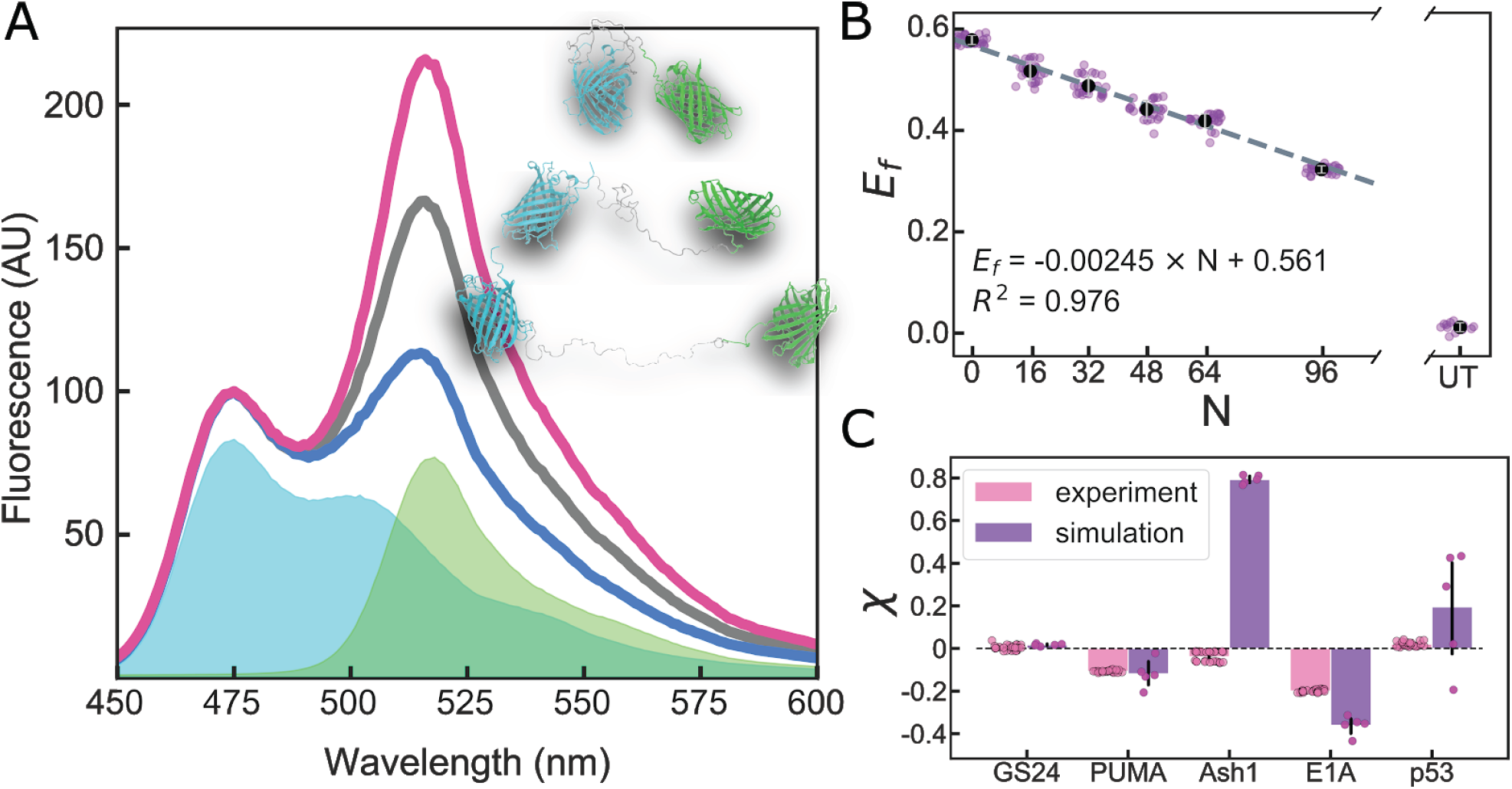
(**A**) Fluorescence spectra normalized to donor peak intensity of a FRET construct in compacting (red), buffer (black), and expanding (blue) solutions. Cyan and green areas are base spectra of donor and acceptor FPs, respectively. Inset shows single configurations for various degrees of expansion. (**B**) FRET efficiency of Gly-Ser repeat linkers vs. number of residues (*N*) in a buffer solution. UT is a solution of untethered, equimolar donor and acceptor. Dashed line shows linear fit of the data. (**C**) Calculated χ for FRET constructs in buffer determined by experiment (average of four repeats with 6 replicates each) and simulation (average of five repeats). Error bars are SD of all replicates/repeats.

The effects of solution chemical changes on protein structure can be likened to a “tug-of-war” between intra-protein interactions and interactions between protein moieties and the surrounding solution. This tug-of-war is a balance that can be shifted by changes to sequence (mutations or post-translational modifications)^15,16^, but also by changes in the physical-chemical composition of the intracellular environment^9,14^. While the sensitivity of IDRs to solution composition has been discussed^13,17,18^, it has not been systematically characterized. Here we set out to systematically evaluate the sensitivity of IDRs to changes in their surrounding environment.

## A high-throughput approach to reveal IDR dimensions using ensemble FRET

We use “solution space” scanning to characterize IDR sensitivity to solution composition changes. This is analogous to “sequence space” scanning, but uses different chemical environments instead of sequence mutations to probe protein behavior. To scan IDRs in solution space at high throughput, we developed a protocol that leverages ensemble FRET to report on changes in the average distance between their termini. We use a protein construct comprising an IDR of interest sandwiched between two fluorescent proteins (FPs) that together form a Förster resonance energy transfer (FRET) pair (**Fig. 1A**). The FPs selected were mTurquoise2^19^ (donor) and mNeonGreen^20^ (acceptor)^21,22^. We chose four IDRs whose ensembles play functional roles: the 61-residue N-terminal transactivation domain of p53 (p53)^4^; the 34-residue BH3 domain of apoptosis regulating protein PUMA (PUMA)^3^; the 83-residue C-terminal domain of the yeast transcription factor Ash1 (Ash1)^23^; the 40-residue N-terminal domain of the adenoviral hub protein E1A (E1A)^24^. For each of these constructs the FRET efficiency,*E*_*f*_, was determined as described in **SI Section S1**.**5**.

To derive changes in IDR dimensions from *E*_*f*_ we began by measuring a series of Gly-Ser (GS) repeats in our FRET backbone to generate a length-dependent point of reference. *E*_*f*_ for these constructs scales linearly with GS repeats as expected (**Fig. 1B**, and see **SI Section S1**.**6**), allowing us to interpolate *E*_*f*_ for a GS linker of a given length to create a ratio χ:

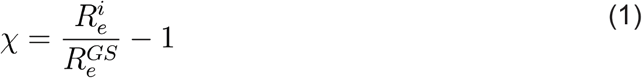

where *R*_*e*_ is the end-to-end distance between donor and acceptor FPs obtained from *E*_*f*_ as described in **SI Section S1**.**7**, and the superscript *i* or *GS* refers to a specific IDR sequence or a GS linker of the equivalent length, respectively. Thus, a negative χ value indicates the chain is more compact, while a positive value indicates it is more expanded, than a GS linker of the equivalent length. Conveniently, χ allows us to plot IDRs of different lengths on the same axes. Our calculations of χ in neat buffer for different FRET constructs reveal a range of behaviors, with Ash1 and p53 having more expanded, and PUMA and E1A more compact, ensembles (**Fig. 1C**).

## IDR ensemble dimensions are sensitive to solution composition and protein sequence, but not to length

We next investigate how IDR dimensions change in different chemical environments. The solutions we use are *not* representative of the cellular environment. Instead, solutions containing osmolytes, polymeric crowders, polyols, free amino acids, denaturants and salts probe IDR structure by “pushing” or “pulling” against the attractions or repulsions of intra-protein interactions. We calculated χ for each combination of IDR/solution as described in **SI Section S1**.**7**. The resulting changes in χ reveal a distinctive solution-space “fingerprint” for each IDR (**Fig. 2A**) and highlight that different sequences have different sensitivities to the same solute^25,26^. This is in sharp contrast to the sensitivity of GS linkers, which all display a similar fingerprint regardless of length (**Fig. S1**).

**Figure 2.**
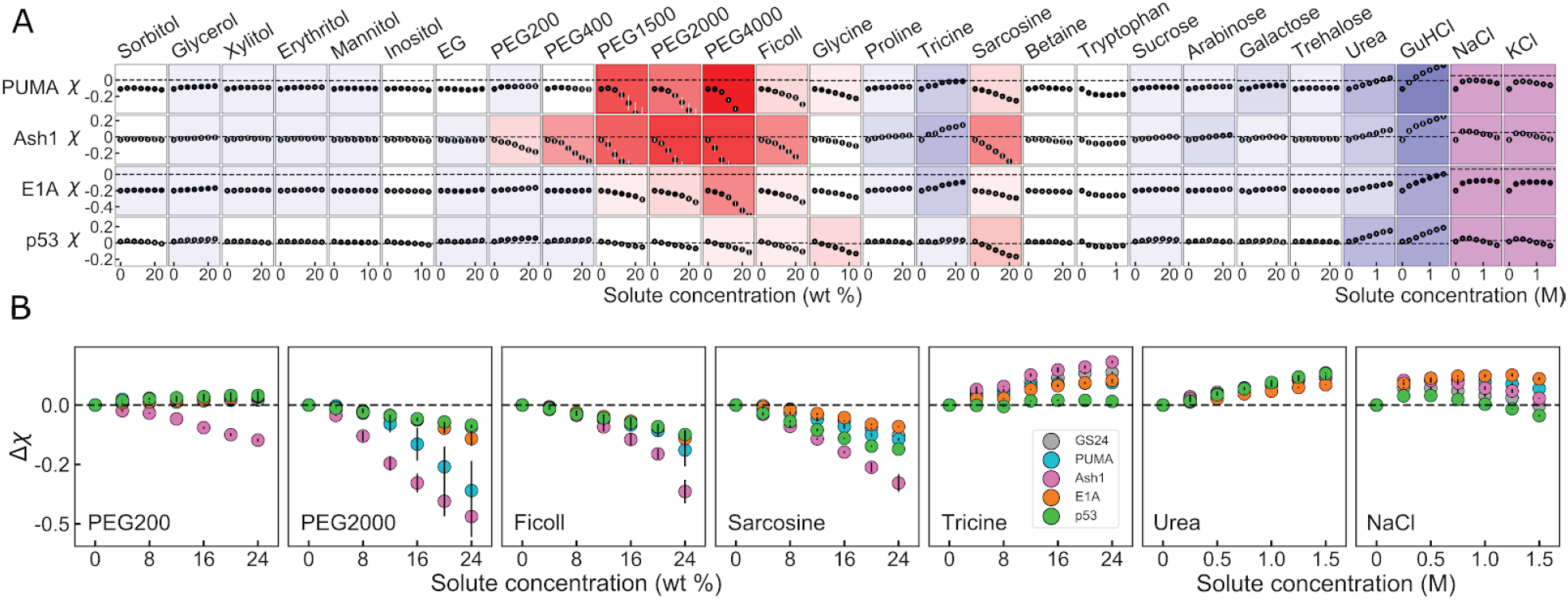
(**A**) Solution space scans of IDRs. Each data point shows the average χ vs. concentration of a specific solute for each protein taken from two repeats. Vertical lines show spread of repeats, and are often too small to see. Proteins vary across columns, and solutes across rows. Background color represents the sensitivity of change to solute addition: stronger colors imply higher sensitivity, red hues indicate compaction, and blue hues indicate expansion. Purple background indicates non-monotonic behavior. (**B**) Differential response of IDRs to individual solutes. Each panel point is the average of the solution-induced change in χ, Δ_χ_, vs. concentration from two repeats of a specific solute for several different constructs. Vertical lines are the spread of the repeats.

Focusing on several solute archetypes reveals interesting trends (**Fig. 2B**). Short polyethylene glycol (PEG) chains, such as PEG200, display disparate effects on different sequences, causing only Ash1 to compact, and the rest to expand, in line with other observations^27^. Larger polymers such as PEG2000 and Ficoll appear to compact the dimensions of all IDRs as shown for other disordered proteins^28^, with a sequence-dependent magnitude that is stronger for Ash1 and PUMA. Smaller solutes like sarcosine and tricine also reveal a linear expanding or compacting effect, but show that different proteins expand or compact by different magnitudes under the same solution. Salts like NaCl display a characteristic non-monotonic effect, as described previously^8,29,30^. In ionic solutes, the initial expansion likely stems from screening of attractive electrostatic interactions that may in fact arise not only from the IDR chain but also from the FP tags, as indicated by the effect on uncharged GS linkers (**Fig. S1**), while the compaction trend stems from specific ion effects, and differs between protein types^31^. Overall, the picture that emerges is that different solution environments affect IDRs in a way that strongly depends on sequence composition and arrangement, but much less on length. The full dataset is available in **Tables S1-S2**.

## IDR dimensions in neat buffer predict sensitivity to solution changes

To see how sensitivities play out in a larger range of IDRs, we turn to all-atom simulations. We use the ABSINTH forcefield that has previously been shown to reproduce experimentally measured IDR ensembles (see **SI Section 2**.**1**).^23,32–34^ To maintain connection with experiments, we start by simulating GS linkers of various lengths (**Fig. S2**), and use ensemble-averaged *R*_*e*_ to calculate χ for simulated IDRs according to **Eq. 1**. The simulation-derived χ for the four different proteins used in our experiments qualitatively agrees with our FRET experiments, aside from Ash1 which is significantly more expanded in simulations than our FRET experiments show (**Fig. 1C**). It is important to note that the absence of FPs in these simulations dictates that the value of χ is necessarily different between experiment and simulations, and a quantitative match is not expected.

We next wanted to see how other naturally occurring disordered sequences would respond to different solution conditions. We have previously designed and calibrated an approach to perform computational solution space scanning with ABSINTH.^12^ We selected 70 experimentally identified IDRs^35^, and used our computational solution space scanning approach to change interactions between the solvent and the backbone of these proteins, akin to the effect of osmolytes and denaturants^36–38^ (data in **Table S3)**. We quantified the sensitivity of the protein to compacting or expanding solutions based on the extent of change in χ (**Fig. S3**). The dataset is sorted from compact to expanded (negative to positive χ) in **Fig. 3A** and shows little correlation with many sequence-based parameters, but relatively strong correlation with the change Δ_χ_ = χ(*solution*) − χ(*buffer*) in solutions that cause the sequence to compact (repulsive solutions) or expand (attractive solutions). We refer to this value as the “solution sensitivity” of the protein. We plot the solution sensitivity, Δ_χ_, vs. protein dimensions in buffer, χ, in **Fig. 3B**. As expected from **Fig. 3A**, sequences with a negative χ have a larger tendency to expand, but a limited ability to compact, and vice versa for positive χ. Remarkably, both compaction and expansion show the same dependence on χ, even at different solution interaction strengths (see **Fig. S4**).

**Figure 3.**
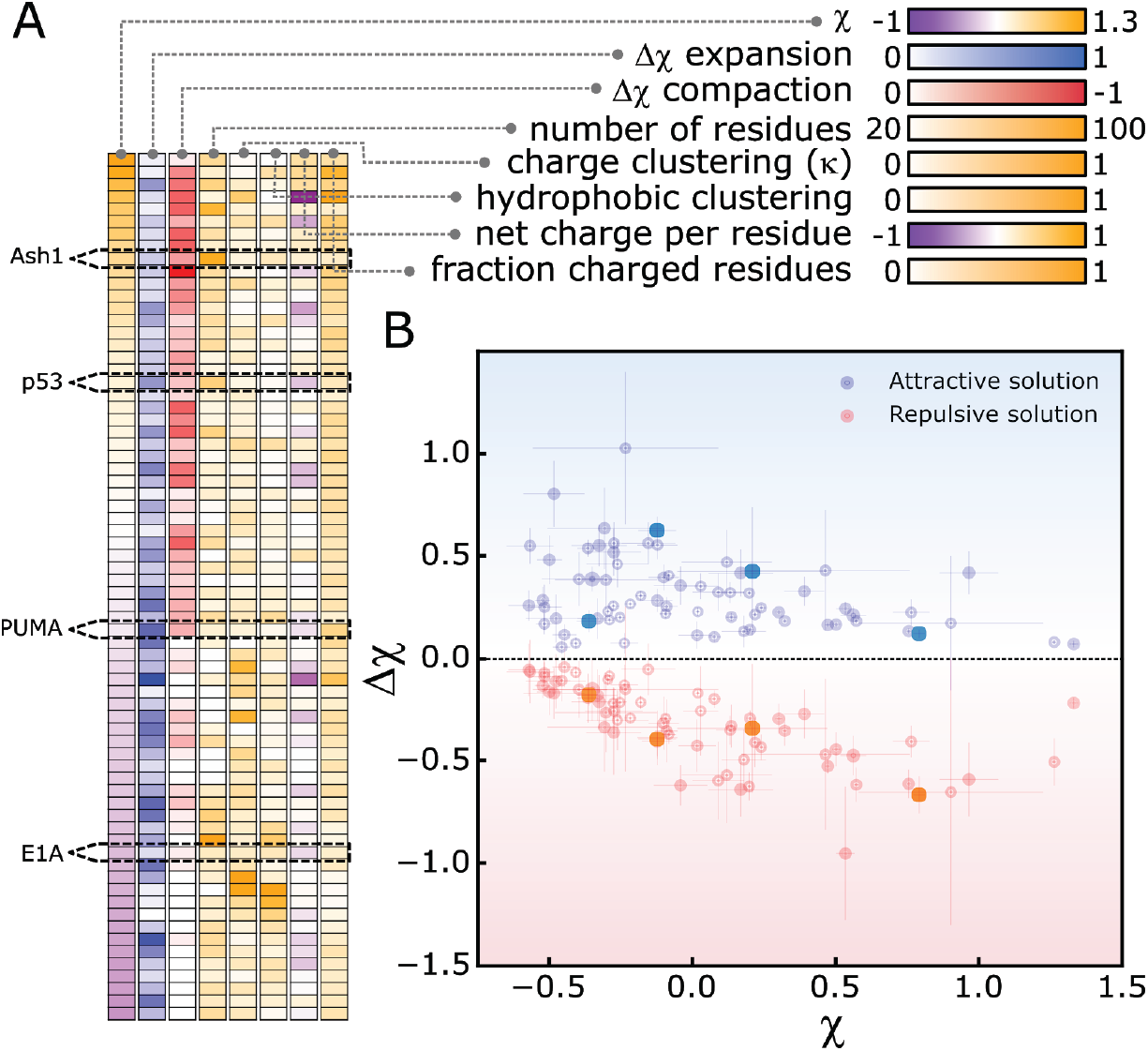
All-atom simulations of IDR sensitivity to solutions. (**A**) Heatmap of protein sensitivity and molecular features. Protein identity varies from top to bottom across cells, and molecular features vary left to right. Colormaps are shown for each molecular feature. (**B**) The magnitude Δ_χ_ in attractive (blue) or repulsive (red) solutions as a function of χ in aqueous solution for each protein in (A). Error bars calculated from SD of 5 repeats. All data available in **Table S3**.

## Predicting the extent of solution sensitivity in intrinsically disordered chains

To see if the non-monotonic trend shown in **Fig. 3B** can be generalized, we measured solution-induced expansion and compaction in a lattice-based heteropolymer model detailed in **SI Section 2**.**2**.^23,34^. We simulated a total of 10^4^ sequences with lengths ranging from 20 to 100 residues in 11 solution conditions, and quantified χ and Δ_χ_ for each sequence/solution pair (**Fig. S5**). The trends from all-atom simulations, re-drawn as a density map in **Fig. 4A**, match the coarse-grained simulations shown in **Fig. 4B**. A non-monotonic change Δ_χ_ in is observed, with the inflection point centered approximately around χ= 0.0 and a ‘dead zone’ in the center of the plot. For naturally expanded chains (χ → 0.4) solution sensitivity is minimized, while for naturally compact chains (χ → –0.4) a broad distribution of sensitivity is observed with respect to expansion, while sensitivity through compaction trends to zero.

**Fig. 4.**
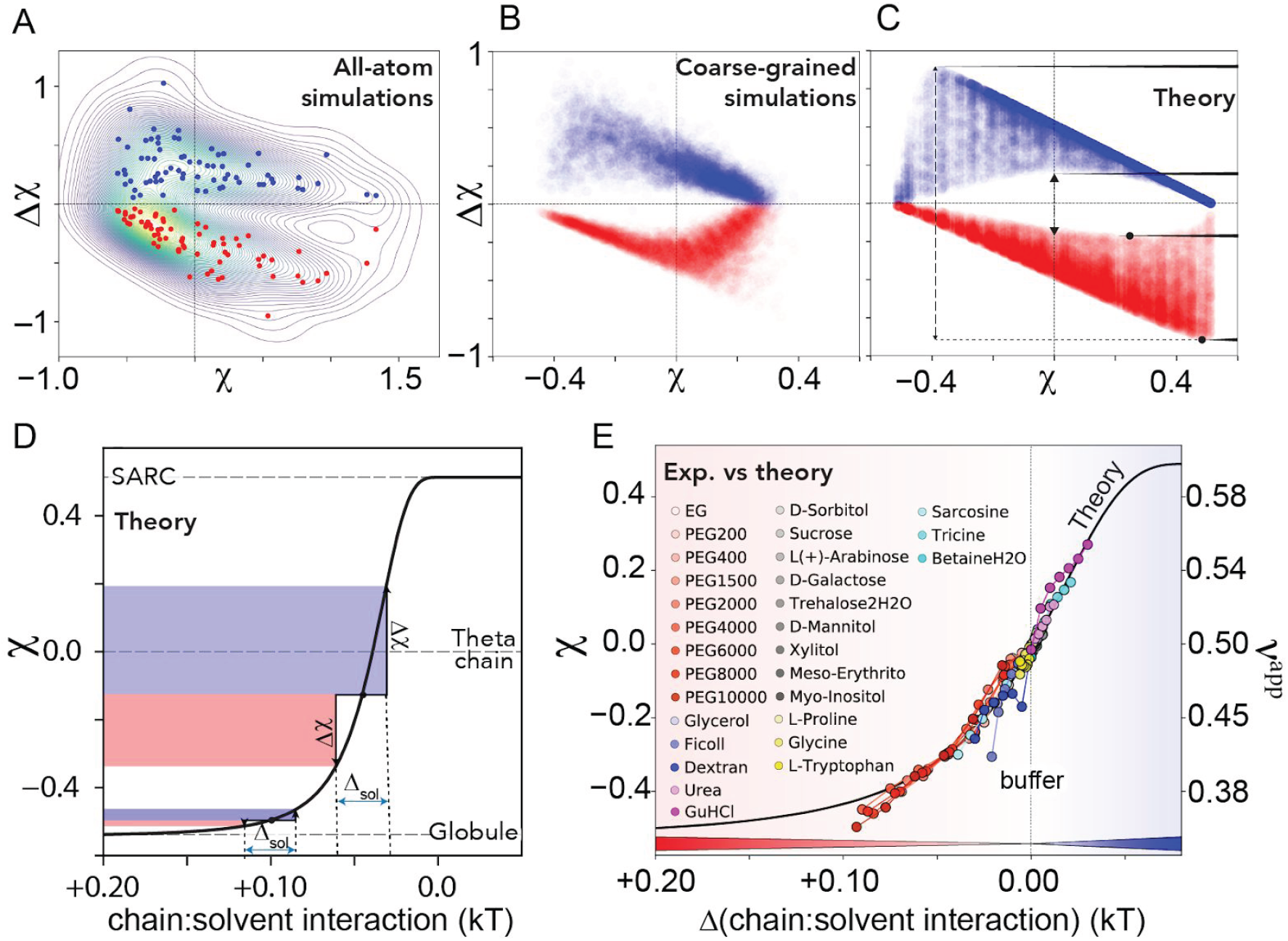
**(A-C).** Density maps of all-atom simulations shown in Fig, 3B (A), PIMMS coarse-grained simulations (B), and an analytical model (C) for solution sensitivity Δ_χ_ vs dimensions in aqueous buffer χ. **(D)** Coil-to-globule transition obtained from an analytical model (SARC = self-avoiding random coil). Δ_χ_ is measured as the height of the blue (contraction) or red (expansion) shaded regions. When the same chain-solvent perturbation (Δ_sol_) is applied to a 100-residue chain with different starting χ values, very different are expected. **(E)** Projection of experimental data for Ash1 onto the analytical model from (D), with solute concentrations scaled to the change in mean-field chain-solvent interaction as compared with neat buffer. Chain dimensions are also shown by their apparent scaling exponent *v*^*app*^. Mapping of other proteins is shown in **Fig. S7**.

Based on these results, we developed an analytical homopolymer model to relate changes in chain-solvent interaction to chain dimensions (see **SI Section 2**.**3**). Using this model we generated chains with a specific χ value in buffer and perturbed the chain-solvent interactions, and directly calculated Δ_χ_ (**Fig. 4C, Fig. S6**). Despite being a simplified homopolymer model, our analytical expression revealed the same phenomenological pattern as obtained in our all-atom and coarse-grained simulations.

The χ-dependence of the chain-solvent interaction strength is shown in **Fig. 4D** (black line), which reveals that Δ_χ_ depends on both the strength of the change in chain-solvent interaction and the χ value in an aqueous solution. Our model offers direct physical intuition as to the origin of the complex relationship between χ and Δχ. Perhaps most importantly, it implies that while expanded or compact proteins display a wide range of sensitivities, IDRs where is around 0 display a basal sensitivity to solution interactions. In this region, where χ most IDRs fall^39^, even small changes to solution composition are predicted to have a measurable effect on IDR dimensions and/or residual structure.^12^

Under the assumption that cosolute-protein interactions scale linearly^25,40^, we globally fit our experimental data onto our analytical model leveraging the fact that all experimental measurements start in the same neat buffer (**Fig. 4E, Fig. S7**). All solution perturbations can be rationally interpreted as driving sequence-dependent shifts along the coil-to-globule transition, in which the magnitude of the shift maps directly to modulation of chain-solvent interactions. The scaling factors required for this mapping qualitatively mirror known co-solute interaction coefficients, and reveal quantitative sequence-dependent differences in the solution response (**Fig. S8**). Chain dimensions can also be represented using an apparent scaling exponent (*v*^app^) (see **SI Section 2**.**4**)^41,42^. The solvent-induced changes observed are substantial, and for many solutes drive changes equal to or greater than changes observed in IDRs due to mutagenesis^34^.

In this work we set out to measure the ability of IDRs to respond to chemical composition changes in their surrounding solution. Although the solutions used here do not represent real cellular environments, they reveal that IDR ensembles carry an inherent, sequence-encoded sensitivity to changes in their chemical environment. This sensitivity can stem from different molecular features, and as far as we determined correlates only with the dimensions of the sequence (χ) in aqueous buffer. IDR function through conformational selection has been reported for numerous proteins. In this mechanism, function is linked to the conformational ensemble of the IDR, directly linking environment-induced ensemble changes to IDR activity. The most exciting idea our data suggests is that changes in the chemical composition which commonly occur in the cell can tune the function (or malfunction) of intrinsically disordered proteins.

## Supporting information

Supplemental Information - Methods, analysis, and figures

Supplemental Table 1 - Raw flourescence emission data for all FRET constructs and FP controls

Supplemental Table 2 - Experimental data used to produce Fig. 2

Supplemental Table 3 - Simulation data for Fig. 3

Supplemental Table 4 - Sequences for all experimental protein constructs

## Acknowledgements

We gratefully acknowledge computing time on the MERCED cluster at UC Merced, funded by NSF Grant No. ACI-1429783, and on the XSEDE computational infrastructure framework, grant No. TG-MCB190103 to ASH and SS, supported by NSF grant No. ACI-1548562. Research reported in this publication was supported by the NIH under award R35GM137926 to SS. DM is supported by a fellowship from NSF-CREST Center for Cellular and Biomolecular Machines at UC Merced Grant No. NSFHRD-1547848.

## References

(1) Wright, P. E.; Dyson, H. J. Intrinsically Disordered Proteins in Cellular Signalling and Regulation. Nat. Rev. Mol. Cell Biol. 2015, 16 (1), 18–29.

(2) Arai, M.; Sugase, K.; Dyson, H. J.; Wright, P. E. Conformational Propensities of Intrinsically Disordered Proteins Influence the Mechanism of Binding and Folding. Proc. Natl. Acad. Sci. U. S. A. 2015, 112 (31), 9614–9619.

(3) Wicky, B. I. M.; Shammas, S. L.; Clarke, J. Affinity of IDPs to Their Targets Is Modulated by Ion-Specific Changes in Kinetics and Residual Structure. Proceedings of the National Academy of Sciences 2017, No. 24, 201705105.

(4) Borcherds, W.; Theillet, F.-X.; Katzer, A.; Finzel, A.; Mishall, K. M.; Powell, A. T.; Wu, H.; Manieri, W.; Dieterich, C.; Selenko, P.; Loewer, A.; Daughdrill, G. W. Disorder and Residual Helicity Alter p53-Mdm2 Binding Affinity and Signaling in Cells. Nat. Chem. Biol. 2014, 10 (12), 1000–1002.

(5) Babu, M. M.; van der Lee, R.; de Groot, N. S.; Gsponer, J. Intrinsically Disordered Proteins: Regulation and Disease. Curr. Opin. Struct. Biol. 2011, 21 (3), 432–440.

(6) Smoyer, C. J.; Jaspersen, S. L. Breaking down the Wall: The Nuclear Envelope during Mitosis. Curr. Opin. Cell Biol. 2014, 1, 1–9.

(7) Stewart, M. P.; Helenius, J.; Toyoda, Y.; Ramanathan, S. P.; Muller, D. J.; Hyman, A. A. Hydrostatic Pressure and the Actomyosin Cortex Drive Mitotic Cell Rounding. Nature 2011, 469 (7329), 226–230.

(8) Vancraenenbroeck, R.; Harel, Y. S.; Zheng, W.; Hofmann, H. Polymer Effects Modulate Binding Affinities in Disordered Proteins. Proc. Natl. Acad. Sci. U. S. A. 2019. https://doi.org/10.1073/pnas.1904997116.

(9) Davis, C. M.; Gruebele, M.; Sukenik, S. How Does Solvation in the Cell Affect Protein Folding and Binding? Curr. Opin. Struct. Biol. 2018, 1, 23–29.

(10) Sukenik, S.; Salam, M.; Wang, Y.; Gruebele, M. In-Cell Titration of Small Solutes Controls Protein Stability and Aggregation. J. Am. Chem. Soc. 2018, 140 (33), 10497–10503.

(11) Tompa, P. The Principle of Conformational Signaling. Chem. Soc. Rev. 2016, 45 (15), 4252–4284.

(12) Holehouse, A. S.; Sukenik, S. Controlling Structural Bias in Intrinsically Disordered Proteins Using Solution Space Scanning. J. Chem. Theory Comput. 2020. https://doi.org/10.1021/acs.jctc.9b00604.

(13) Banks, A.; Qin, S.; Weiss, K. L.; Stanley, C. B.; Zhou, H. X. Intrinsically Disordered Protein Exhibits Both Compaction and Expansion under Macromolecular Crowding. Biophys. J. 2018, 114 (5), 1067–1079.

(14) Hsu, P. P.; Sabatini, D. M. Cancer Cell Metabolism: Warburg and beyond. Cell 2008, 134 (5), 703–707.

(15) Wu, D.; Zhou, H.-X. Designed Mutations Alter the Binding Pathways of an Intrinsically Disordered Protein. Sci. Rep. 2019, 9 (1), 6172.

(16) Bah, A.; Forman-Kay, J. D. Modulation of Intrinsically Disordered Protein Function by Post-Translational Modifications. J. Biol. Chem. 2016, 291 (13), 6696–6705.

(17) van der Lee, R.; Buljan, M.; Lang, B.; Weatheritt, R. J.; Daughdrill, G. W.; Dunker, A. K.; Fuxreiter, M.; Gough, J.; Gsponer, J.; Jones, D. T.; Kim, P. M.; Kriwacki, R. W.; Oldfield, C. J.; Pappu, R. V.; Tompa, P.; Uversky, V. N.; Wright, P. E.; Babu, M. M. Classification of Intrinsically Disordered Regions and Proteins. Chem. Rev. 2014, 114 (13), 6589–6631.

(18) Mansouri, A. L.; Grese, L. N.; Rowe, E. L.; Pino, J. C.; Chennubhotla, S. C.; Ramanathan, A.; O’Neill, H. M.; Berthelier, V.; Stanley, C. B. Folding Propensity of Intrinsically Disordered Proteins by Osmotic Stress. Mol. Biosyst. 2016, 12 (12), 3695–3701.

(19) Goedhart, J.; von Stetten, D.; Noirclerc-Savoye, M.; Lelimousin, M.; Joosen, L.; Hink, M. A.; van Weeren, L.; Gadella, T. W. J., Jr; Royant, A. Structure-Guided Evolution of Cyan Fluorescent Proteins towards a Quantum Yield of 93%. Nat. Commun. 2012, 1, 751.

(20) Shaner, N. C.; Lambert, G. G.; Chammas, A.; Ni, Y.; Cranfill, P. J.; Baird, M. A.; Sell, B. R.; Allen, J. R.; Day, R. N.; Israelsson, M.; Davidson, M. W.; Wang, J. A Bright Monomeric Green Fluorescent Protein Derived from Branchiostoma Lanceolatum. Nat. Methods 2013, 10 (5), 407–409.

(21) Mastop, M.; Bindels, D. S.; Shaner, N. C.; Postma, M.; Gadella, T. W. J., Jr; Goedhart, J. Characterization of a Spectrally Diverse Set of Fluorescent Proteins as FRET Acceptors for mTurquoise2. Sci. Rep. 2017, 7 (1), 11999.

(22) Sørensen, C. S.; Kjaergaard, M. Effective Concentrations Enforced by Intrinsically Disordered Linkers Are Governed by Polymer Physics. Proc. Natl. Acad. Sci. U. S. A. 2019, 116 (46), 23124–23131.

(23) Martin, E. W.; Holehouse, A. S.; Grace, C. R.; Hughes, A.; Pappu, R. V.; Mittag, T. Sequence Determinants of the Conformational Properties of an Intrinsically Disordered Protein Prior to and upon Multisite Phosphorylation. J. Am. Chem. Soc. 2016, 138 (47), 15323–15335.

(24) Ferreon, A. C. M.; Ferreon, J. C.; Wright, P. E.; Deniz, A. A. Modulation of Allostery by Protein Intrinsic Disorder. Nature 2013, 498 (7454), 390–394.

(25) Sukenik, S.; Sapir, L.; Gilman-Politi, R.; Harries, D. Diversity in the Mechanisms of Cosolute Action on Biomolecular Processes. Faraday Discuss. 2013, 1, 225–237; discussion 311–327.

(26) Senske, M.; Törk, L.; Born, B.; Havenith, M.; Herrmann, C.; Ebbinghaus, S. Protein Stabilization by Macromolecular Crowding through Enthalpy rather than Entropy. J. Am. Chem. Soc. 2014, 136 (25), 9036–9041.

(27) Kozer, N.; Kuttner, Y. Y.; Haran, G.; Schreiber, G. Protein-Protein Association in Polymer Solutions : From Dilute to Semidilute to Concentrated. Biophys. J. 2007, 92 (6), 2139–2149.

(28) Zosel, F.; Soranno, A.; Buholzer, K. J.; Nettels, D.; Schuler, B. Depletion Interactions Modulate the Binding between Disordered Proteins in Crowded Environments. Proc. Natl. Acad. Sci. U. S. A. 2020, 117 (24), 13480–13489.

(29) Pegram, L. M.; Wendorff, T.; Erdmann, R.; Shkel, I.; Bellissimo, D.; Felitsky, D. J.; Record, M. T., Jr. Why Hofmeister Effects of Many Salts Favor Protein Folding but Not DNA Helix Formation. Proc. Natl. Acad. Sci. U. S. A. 2010, 107 (17), 7716–7721.

(30) Sarkar, M.; Li, C.; Pielak, G. J. Soft Interactions and Crowding. Biophys. Rev. 2013, 5 (2), 187–194.

(31) Pegram, L. M.; Record, M. T., Jr. Thermodynamic Origin of Hofmeister Ion Effects. J. Phys. Chem. B 2008, 112 (31), 9428–9436.

(32) Vitalis, A.; Pappu, R. V. ABSINTH: A New Continuum Solvation Model for Simulations of Polypeptides in Aqueous Solutions. J. Comput. Chem. 2009, 30 (5), 673–699.

(33) Das, R. K.; Huang, Y.; Phillips, A. H.; Kriwacki, R. W.; Pappu, R. V. Cryptic Sequence Features within the Disordered Protein p27Kip1 Regulate Cell Cycle Signaling. Proc. Natl. Acad. Sci. U. S. A. 2016, 113 (20), 5616–5621.

(34) Martin, E. W.; Holehouse, A. S.; Peran, I.; Farag, M.; Incicco, J. J.; Bremer, A.; Grace, C. R.; Soranno, A.; Pappu, R. V.; Mittag, T. Valence and Patterning of Aromatic Residues Determine the Phase Behavior of Prion-like Domains. Science 2020, 367 (6478), 694–699.

(35) Piovesan, D.; Tabaro, F.; Micetic, I.; Necci, M.; Quaglia, F.; Oldfield, C. J.; Aspromonte, M. C.; Davey, N. E.; Davidović, R.; Dosztányi, Z.; Elofsson, A.; Gasparini, A.; Hatos, A.; Kajava, A. V.; Kalmar, L.; Leonardi, E.; Lazar, T.; Macedo-Ribeiro, S.; Macossay-Castillo, M.; Meszaros, A.; Minervini, G.; Murvai, N.; Pujols, J.; Roche, D. B.; Salladini, E.; Schad, E.; Schramm, A.; Szabo, B.; Tantos, A.; Tonello, F.; Tsirigos, K. D.; Veljković, N.; Ventura, S.; Vranken, W.; Warholm, P.; Uversky, V. N.; Dunker, A. K.; Longhi, S.; Tompa, P.; Tosatto, S. C. E. DisProt 7.0: A Major Update of the Database of Disordered Proteins. Nucleic Acids Res. 2017, 45 (D1), D219–D227.

(36) Liu, Y.; Bolen, D. W. The Peptide Backbone Plays a Dominant Role in Protein Stabilization by Naturally Occurring Osmolytes. Biochemistry 1995, 34 (39), 12884–12891.

(37) Auton, M.; Holthauzen, L. M. F.; Bolen, D. W. Anatomy of Energetic Changes Accompanying Urea-Induced Protein Denaturation. Proc. Natl. Acad. Sci. U. S. A. 2007, 104 (39), 15317–15322.

(38) O’Brien, E. P.; Ziv, G.; Haran, G.; Brooks, B. R.; Thirumalai, D. Effects of Denaturants and Osmolytes on Proteins Are Accurately Predicted by the Molecular Transfer Model. Proc. Natl. Acad. Sci. U. S. A. 2008, 105 (36), 13403–13408.

(39) Marsh, J. A.; Forman-Kay, J. D. Sequence Determinants of Compaction in Intrinsically Disordered Proteins. Biophys. J. 2010, 98 (10), 2383–2390.

(40) Harries, D.; Rosgen, J. A Practical Guide on How Osmolytes Modulate Macromolecular Properties. Methods Cell Biol. 2008, 84 (07), 679–735.

(41) Holehouse, A. S.; Pappu, R. V. Collapse Transitions of Proteins and the Interplay Among Backbone, Sidechain, and Solvent Interactions. Annu. Rev. Biophys. 2018, 1, 19–39.

(42) Riback, J. A.; Bowman, M. A.; Zmyslowski, A. M.; Knoverek, C. R.; Jumper, J. M.; Hinshaw, J. R.; Kaye, E. B.; Freed, K. F.; Clark, P. L.; Sosnick, T. R. Innovative Scattering Analysis Shows That Hydrophobic Disordered Proteins Are Expanded in Water. Science 2017, 358 (6360), 238–241.

