## Supplemental Information - Methods, analysis, and figures for "Probing the Hidden Sensitivity of Intrinsically Disordered Proteins to their Chemical Environment"

### S1 Experimental Methods

##### S1.1 FRET construct design and cloning

FRET backbone (called fIDR_pET-28a(+)-TEV, **Fig. S9**), was prepared by ligating mTurquoise2 and mNeonGreen into pET28a-TEV backbone using 5' NdeI and 3' XhoI restriction sites. Genes encoding for IDR regions were obtained from GenScript, and ligated between the two fluorescent proteins using 5' NdeI and 3' HindIII restriction sites. Cloned plasmids were amplified in XL1 Blue (Invitrogen) cell lines using manufacturer supplied protocol. Sequences of all IDR sequence inserts are available in **Table S4**.

##### S1.2 FRET construct expression and purification

Plasmids encoding for FRET constructs were expressed in BL21(DE3) cells in LB medium with 50 µg/mL kanamycin. Cultures were incubated at 37 °C while shaking at 225 rpm until OD600 of 0.6 was reached (approx. 3 h), then induced with 1 mM IPTG and incubated for 20 h at 16 °C while shaking at 225 rpm. Cells were harvested by centrifugation for 15 min at 3,000 rcf , the supernatant was discarded, and the cells were lysed in lysis buffer (50 mM $NaH_2PO_4$, pH 8, 0.5 M NaCl) using an Avestin Emulsiflex C3 homogenizer. Lysate was centrifuged for 1 h at 20,000 rcf and the supernatant collected and flowed through a column packed with Ni-NTA beads (Qiagen). FRET construct was eluted with 50 mM $NaH_2PO_4$, pH 8, 0.5 M NaCl, 250 mM imidazole, and further purified using size-exclusion chromatography on a Superdex 200 pg column (GE Healthcare) in an AKTA go protein purification system (GE Healthcare). The purified FRET constructs were aliquoted into 200 µL aliquots, flash frozen in liquid nitrogen, and stored at -80 °C in 20 mM sodium phosphate buffer, pH 7.4, with the addition of 100 mM NaCl. Protein concentration was measured after thawing and before use using UV-vis absorbance at 434 and 506 nm (the peak absorbance wavelengths for mTurquoise2 and mNeonGreen, respectively; the molar absorbance coefficients for mTurquoise2 and mNeonGreen are 30,000 $cm^{-1}M^{-1}$ and 116,000 $cm^{-1}M^{-1}$, respectively.[1] Calculations of concentration based on $\lambda$ = 434 nm produced slightly higher values than calculations based on $\lambda$ = 506 nm, so the concentrations based on the measurement at $\lambda$ = 506 nm were used), and purity was assessed by SDS-PAGE after thawing and before use.

##### S1.3 Solution preparation and specifics

Solutes were purchased from Alfa Aesar (Dextran, Xylitol, L-Tryptophan, Sarcosine, PEG200, PEG400, PEG1500, PEG2000, PEG4000, PEG6000, PEG8000, PEG10000), VWR (D-Sorbitol), GE Healthcare (Ficoll), TCI (Meso-Erythritol, D-(+)-Trehalose Dihydrate), Thermo Scientific (Guanidine Hydrochloride), Acros Organics (D-Mannitol, Betaine Monohydrate, L(+)-Arabinose), Sigma-Aldrich (Myo-Inositol, Taurine), and Fisher BioReagents (Ethylene Glycol, D-Galactose, Glycerol, Glycine, L-Proline, Tricine, Potassium Chloride, Sodium Chloride, Urea), and used without further purification. Stock solutions were made by mixing the solute with 20 mM sodium phosphate buffer, pH 7.4, with the addition of 100 mM NaCl except for NaCl and KCl solutions, which were free of additional salt. The same buffer was used for all dilutions.

##### S1.4 FRET experiments

FRET experiments were conducted in black plastic 96-well plates (Nunc) using a CLARIOstar plate reader (BMG LABTECH). Buffer, stock solution and purified protein solution were mixed in each well so as to reach a volume of 150 µL containing the desired concentrations of the solute and the FRET construct, with a final concentration of 1 µM protein (or of each FP in the case of the "untethered" control). Fluorescence measurements were taken from above, at a focal height of 5.7 mm, with gain fixed at 1020 for all samples. For each FRET construct, two repeats with 12 replicates each were performed for each protein in neat buffer, and at least two repeats were done in every other solution condition. Fluorescence spectra were obtained for each FRET construct in each solution condition by exciting the sample in a 16-nm band centered at $\lambda$ = 420 nm, with a dichroic at $\lambda$ = 436.5 nm, and measuring fluorescence emission from $\lambda$ = 450 to 600 nm, averaging over a 10 nm window moved at intervals of 0.5 nm. Base donor and acceptor spectra for each solution condition were obtained using the same excitation and emission parameters on solutions containing 1 µM mTurquoise2 or mNeonGreen alone, and measuring fluorescence emission from 450 to 600 nm[1,2].

##### S1.5 Calculation of $E_f$

The process of calculating the FRET efficiency $E_f$ for a FRET construct in one solute at a range of concentrations is summarized in **Fig. S10**. Specifically, $E_f$ of each FRET construct in each solution condition was calculated by linear regression of the fluorescence spectrum of the FRET construct with the spectra of the separate donor and acceptor emission spectra (in order to correct for solute-dependent effects on fluorophore emission) in the same solution conditions. $E_f$ was calculated using :

$$E_f = 1 - \frac{F_d}{\frac{Q_d f_d}{Q_a f_a} F_s + F_d}$$

where $F_d$ is the decoupled donor contribution, $F_a$ is the decoupled acceptor contribution, $f_d$ is the area-normalized donor spectrum, $f_a$ is the area-normalized acceptor spectrum, $Q_d = 0.93$ is the quantum yield of the donor, and $Q_a = 0.8$ is the quantum yield of the acceptor[2,3].

More specifically, the data for each series of solution conditions consisting of increasing concentrations of a single solute was processed in the following manner:

1. Raw spectra for the free donor and free acceptor in the various solution conditions were loaded, and the averages of all repeats in each solution condition were computed. These averages are referred to as the "raw" donor and acceptor spectra below because they will be further corrected.

2. The donor and acceptor peak intensities were assumed to change in a linear fashion with increasing solute concentration, so peak height of donor or acceptor-only spectra *vs.* concentrations were linearly fit.

3. To correct for artifacts (such as variations in FRET construct concentration between different wells) that may contribute to unexpected differences in fluorescence intensity, a correction factor was applied to each raw donor and acceptor spectrum to bring the peak intensity to the linear fit described in step 2, resulting in "corrected" donor and acceptor spectra. Importantly, while this corrected well-to-well variations in raw data, it did little to affect the overall values or trends in $\chi$ (e.g., without this correction Fig. 1 and 2 would vary by less than 5%).

4. The raw FRET construct fluorescence spectra for the series were loaded.

5. To compensate for unintended direct excitation of the acceptor by donor excitation frequency, the corrected acceptor spectrum for each solution condition was subtracted from the FRET construct spectrum for each solution condition, resulting in "corrected" FRET construct spectra.

6. The corrected donor, acceptor and FRET construct spectrum for each solution condition was fitted with a linear regression function to determine the decoupled contributions of the donor and acceptor to the FRET construct spectrum.

7. $E_f$ of each FRET construct in each solution condition was calculated using the equation shown above.

##### S1.6 Assessment of the expected scaling behaviour for interprotein distances

For flexible polymers, the end-to-end distance ($R_e$) and radius of gyration ($R_g$) follow well-defined scaling relationships defined by $R = AN^{\nu}$ where $R$ is a physical distance (i.e., $R_e$ or $R_g$), $A$ is a constant in units of distance, $N$ is the unitless degree of polymerization (i.e., number of residues) and $\nu$ is a unitless apparent scaling exponent [4,5]. For constructs that have two folded domains connected by a flexible linker, in the limit of infinitely long linkers the inter-fluorescent protein distance will approximately equal the end-to-end distance of the intervening linker. However, in the limit of finite-length linkers where the linker dimensions are on a par with the dimensions of the folded proteins, we anticipated that deviations from conventional scaling theory may arise at least in part due to the excluded volume effects of the fluorescent proteins.

To assess the excluded volume component of this deviation, we examined the expected intra-fluorescent protein distance dependence on linker length for a well-defined self-avoiding random coil system. Such a model is convenient in that the dependence of the end-to-end distance for a flexible self-avoiding polymer is well defined analytically as $R_e = BN^{0.59}$.

We built a series of fluorescent-protein linker constructs with linkers of various lengths and performed simulations at all-atom resolution using the CAMPARI simulation engine and the ABSINTH implicit solvent model (see also **SI Section S2.1**). To achieve behavior in the true self-avoiding random coil limit, the Hamiltonian (which here refers to the instantaneous potential energy function) used to generate the ensemble does not experience a contribution from the attractive portion of the Lennard-Jones potential for short-range non-bonded interactions, nor solvation effects, nor electrostatic interactions, as described previously[6]. The backbone dihedral angles on the two fluorescent proteins were held fixed while the backbone dihedral angles of the linker were allowed to vary. All side chains were fully flexible. In effect, this provides a “toy” system in a well-defined polymer limit which allows us to assess the impact of the fluorescent proteins without any confounding concerns for forcefield accuracy, sampling challenges, etc.

We first established that a flexible linker between two FPs indeed scales as expected for a self-avoiding random coil. The scaling exponent obtained by fitting a number of GS repeats vs. inter-GS distance revealed a scaling exponent of 0.61 - extremely close to the value of 0.59 expected from analytical theory (**Fig. S11**)

We then repeated the same analysis for the same system assessing the intra-domain distance between the chromophores in the fluorescent proteins - i.e., the inter-fluorescent protein distance (**Fig. S12**). Unlike the inter-chain distance obtained, here we obtained a linear dependence of inter-fluorescent protein distance on GS length. This behavior is readily explained by the excluded volume impact of the fluorescent proteins. For shorter chains the inter-fluorescent protein distance is much larger than an analogous flexible polymer because the excluded volume from the fluorescent proteins effectively act as repulsors at the chain ends. However, as chain length increases this effect becomes less significant, the offset becomes negligible and the system returns to a power-law dependence. This behavior is not specific to the self-avoiding random coil, and as such we expected an approximately linear dependence of inferred distance on the number of GS repeats. Indeed, this linear dependence mirrors what we observed experimentally, providing confidence that our experimentally-derived distances are following expected trends given the physical nature of the setup.

##### *S1.7 Calculation of* $\chi$

For each FRET construct in each solution condition, $\chi$ was calculated in three steps:

1. The mean FRET efficiency values for 24 replicates (in 4 repeats) each for linkers of 8, 16, 24, 32 and 48 GS repeats (16, 32, 48, 64 and 96 amino acids in length) in a buffer solution (20 mM $NaH_2PO_4$, 100 mM NaCl) were linearly fit to arrive at a relation between FRET efficiency in buffer to the number of amino acids (N) in the GS linker.

2. That slope and y-intercept (shown in **Fig. 1B**) were used to interpolate an implied FRET efficiency ($E_f^{GS}$) for a GS linker of the same length N as the IDR of interest.

3. $\chi$ was then calculated as:

$$\chi = \frac{R_{ee}^i}{R_{ee}^{GS}} - 1 = \frac{R_0^i(1/E_f^i - 1)^{\frac{1}{6}}}{R_0^{GS}(1/E_f - 1)^{\frac{1}{6}}} - 1 = \frac{n^i(1/E_f - 1)^{\frac{1}{6}}}{n^{GS}(1/E_f^{GS} - 1)^{\frac{1}{6}}} - 1$$

where $R_0$ is the Förster distance, defined as the distance between the FPs at which $E_f$ = 0.5, the superscript $i$ indicates the protein we are measuring, the superscript $GS$ indicates a GS linker of length equivalent to that of protein $i$, and $n$ is the refractive index of the solution in which the protein is measured. We have tried modulating the refractive index between 1.33 (for neat buffer) and 1.37 (the refractive index of 24 w/w% PEG10000)[7] and noticed no significant changes in the trends of our data, and an absolute change of < 5% in absolute values of $\chi$. We therefore decided not to use this correction for the work presented in **Figs. 1** and **2**.

### S2 Computational Methods

##### S2.1 All-atom simulations

All-atom Monte Carlo-based simulations were performed using the CAMPARI simulation suite, with the ABSINTH implicit solvent model[8]. In CAMPARI, the effective Hamiltonian is a combination of 4 energy terms:

$$E_{total} = W_{solv} + U_{LJ} + W_{el} + U_{corr}$$

Here $U_{LJ}$ is the Lennard-Jones potential between protein residues, $W_{el}$ is the electric potential term based on coulombic potential, $U_{corr}$ is a term applied to the dihedral angles, and $W_{solv}$ is a solution-protein interaction term based on the ABSINTH implicit solvent model[9], which is equivalent to a transfer free energy from a vacuum to a dilute aqueous solution.

Our solution space scanning method is carried out as described previously[10]. Briefly, the implicit solvation term, $W_{solv}$, is first calculated for each sequence based on its fully extended protein conformation. This represents the maximum transfer free energy ($W_{solv}^{max}$) since it is the most exposed configuration accessible to the protein. Solution space is then probed by modulating $W_{solv}^{max}$ by changing the attraction/repulsion of different protein moieties in relation to the implicit solvent. We express the total strength of solution interaction by the parameter $\psi$ where

$$\psi = \frac{W_{solv}^{max}(solution) - W_{solv}^{max}(water)}{W_{solv}^{max}(water)} \times 100\%$$

In this paper we change $\psi$ by making interactions with the backbone less or more attractive (negative or positive $\psi$ values, respectively). Previous calibration based on helix to coil transition has shown that a 1 M Urea solution is equivalent to $\psi \approx +1.2\%$.[10]

Our all-atom simulation dataset consists of 70 proteins (not including GS repeats). We selected sequences that were shown experimentally to be disordered, as collected on the DisProt server[11]. All sequences were simulated at 310 K with $10^7$ steps of equilibration, followed by $7\times10^7$ steps of production. Conformations were written every 12,500 steps, resulting in a total of ~ 5,000 conformations for every simulation. Each sequence was simulated in 5 independent repeats, resulting in an ensemble containing a total of 20,000 conformations per sequence. The MDtraj python library[12] was used to calculate the radius of gyration and end-to-end distance of the ensemble. Data from analyzed all-atom trajectories for each sequence is available in **Table S3**.

##### S2.2 Coarse-grained simulations

Our coarse-grained depiction of heteropolymer IDPs uses the PIMMS simulation framework[13]. PIMMS is a lattice-based Monte Carlo simulation engine in which inter-bead interactions are determined by nearest-neighbor interactions. All bead interactions are anisotropic along on-lattice and diagonal directions. The system evolves through a collection of moves that include individual crankshaft moves, chain translation/rotation, and chain pivot moves. For our purposes, residues are represented as beads, and a simple heteropolymer amino acid alphabet was used to generate chains of various lengths with a heteropolymeric distribution of residues that are similar to polar, hydrophobic, and charged amino acid residues. We emphasize that the parameters generated here, shown in **Fig. S5A**, are phenomenological and not meant to reflect specific amino acids. The set of PIMMS sequences used are available upon request.

The parameters chosen demonstrate sequence-specific coil-to-globule transitions, as shown in **Fig. S5B**. The simulation temperature was set to be units of $k_B T$. Accordingly, the total energy of the system in a given state is calculated based on a summation over pairwise interactions involving nearest-neighbor, non-bonded contacts, or a solution interaction in the case that no neighbors are present. Moves are accepted or rejected via a standard Metropolis criterion whereby the acceptance ratio is $\min\{1, \exp(-\Delta E/k_B T)\}$ where $k_B = 1$, $\Delta E$ is the energy difference between the current and proposed configurations, and T has the same units as the contact energies thus making the ratio $\Delta E/k_B T$ a dimensionless quantity. This conversion makes the point that the parameterized interactions that reproduce the observed experimental data are in fact relatively weak, being less than $k_B T$, depending of course on the simulation temperature.

For each chain length, 2000 sequences were randomly generated, and each sequence simulated in 10 solution interaction strengths (plus "buffer" condition) for a total of 11 trajectories per sequence. Each simulation consisted of a 20-step equilibration followed by a 1020-step production run at T=70. Sequence analysis was printed out every 10 steps, and the reported distances are averages of the entire trajectory. Simulations were performed in a sufficiently large box to avoid finite size effects. Average end-to-end distances vs. solution interaction strengths for the entire dataset are shown in **Fig. S5B**.

##### S2.3 Analytical model for the solution-driven coil-to-globule transition of a polymer

We developed a simple and generic analytical model to characterize the coil-to-globule transition of a homopolymer as assessed by a mean-field net inter-monomer interaction parameter. This model was then parameterized using homopolymeric PIMMS simulations performed for a range of chain lengths and interaction strengths to provide an analytical expression that relates the inter-monomer interaction strength to the degree of compaction/expansion as measured by the parameter $\chi$. While we "parameterize" using PIMMS simulations, the simulations essentially tailor the model parameters to reproduce the interaction strengths and dimensions as are native to PIMMS. In principle any polymer model could be used to obtain key numerical parameters that dictate spatial and interaction features.

This model is built on the assumption that the coil-to-globule transition can be empirically mapped as a cooperative transition in which the cooperativity and midpoint show an exponential dependence on chain length, and the end-points reflect defined expected $\chi$ values for a flexible polymer in the globule (compact) or coil (expanded) limits. Specifically, we define $\chi$ as

$$\chi = a + b\left(\frac{1}{(m/e)^{\theta}}\right),$$

Where

$$\theta = c\log(L) + d$$

and

$$m = \gamma L^{\gamma}$$

$$a = \left(\frac{L^{0.33}}{L^{0.50}}\right) - 1$$

$$b = \left[\left(\frac{L^{0.59}}{L^{0.50}}\right) - 1\right] - a$$

The parameters in this model are defined as follows:

- $L$ is chain length
- $e$ is the apparent net inter-monomer interaction energy (measured in kT)
- $\theta$ is a measure of the cooperativity of the coil-to-globule transition, and itself depends logarithmically on $L$ and two free parameters ($c$ and $d$)
- $m$ is a measure of the midpoint of the coil-to-globule transition, and depends exponentially on chain length and one free parameter ($\gamma$)

The free parameters ($c$, $d$, and $\gamma$) are obtained by fitting to homopolymeric PIMMS simulations where $\chi$ is calculated directly from the simulations (**Fig. S13-S14**). The specific values for these three parameters will depend on the physical nature of the polymer model, but do not ultimately influence the limiting behavior or trends of the model behavior assuming they retain physically realistic values. These parameters depend on chain stiffness and monomer valence.

This model was chosen to provide a simple analytical description, under the simplifying assumption that chain solvent-dependence can, to a first-order approximation, be described using a simple homopolymer that expands/compacts as reported by $\chi$. Chain-solvent interactions are captured in terms of an apparent intra-bead interaction parameter (*e*) which reports on the net favorable energy associated with monomer-monomer interaction in a given solution.

In the limit of a self-avoiding chain, the coil-to-globule transition is entirely determined by the chain-solvent interaction strength. In the limit of a chain where chain-solvent interactions are set to zero, the coil-to-globule transition is entirely determined by the monomer-monomer interaction strength. Real chains sit somewhere between these two limits, where both chain-chain and chain-solvent interactions contribute to the chain dimensions. Our model is formally parameterized in the non-interacting chain-solvent limit, but this can be recast as the non-interacting chain-chain limit in which the apparent chain-solvent interactions are defined as half the chain-chain interactions. In this way, we can write the coil-to-globule transition as either a function of chain-chain interactions or chain-solvent interactions, as is shown in **Fig. S15.** For simplicity we have leveraged the chain-solvent representation, as most easily dovetails with our experimental work.

In its current format the maximum chain expansion reflects the self-avoiding chain limit in which chain-solvent interactions are set to zero. Note that for polypeptides with charged residues, further expansion is possible via electrostatic repulsion.[14] These longer-range repulsive interactions are not captured by our analytical model nor by the model parameters used for our PIMMS simulations. However, they are evident in our all-atom simulations, offering an explanation as to why the $\chi$ axes for the all-atom simulations extend to substantially larger values than in either the theory or coarse-grained simulations.

**S2.4 Converting from χ to ν**

As in **Eq. 1** we define $\chi$ as

$$\chi_i = \frac{R_e^i}{R_e^{GS}} - 1$$

Considering $R_e$ can also be written as

$$R_e^i = BN^{\nu_i^{app}}$$

Where $B$ is a prefactor in units of distance and the apparent scaling exponent ($\nu$) is a measure of the apparent solvent quality for the chain[4,15]. In both our simulations and prior experiments, a GS linker in neat buffer behaves as a polymer in a theta solvent, a reference state in which chain-chain and chain-solvent interactions are counterbalanced, and where $\nu = 0.5$[16]. Operating under this assumption, we can rewrite $\chi$ as

$$\chi_i = \frac{BN^{\nu_i^{app}}}{BN^{0.5}} - 1$$

And more simply as

$$\chi = \frac{N^{\nu_i^{app}}}{N^{0.5}} - 1$$

As such, it is trivial to convert between $\chi$ and and the apparent scaling exponent ($\nu$) for the chain of a given length $N$ in the limit of a homopolymer instantiation of our model under the simplifying assumption of a fixed, sequence-independent and $\nu$-independent prefactor ($B$). For heteropolymers this assumption may not be valid, but as applied to our simple homopolymer model this is a reasonable set of approximations.

### Supplementary Figures

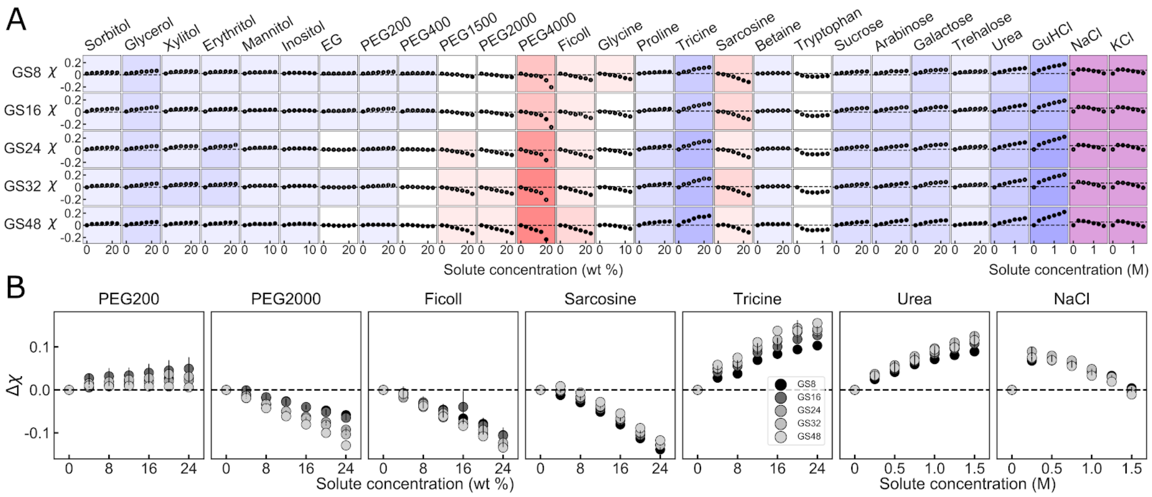


**Figure S1.** (**A**) Solution space scans of Gly-Ser linkers. Each data point shows the average $\chi$ vs. concentration of a specific solute for each protein taken from two repeats. Vertical lines show spread of repeats, and are often too small to see. Proteins vary down columns, and solutes vary along rows. Background color represents the sensitivity of change to solute addition: stronger colors imply higher sensitivity, red hues indicate compaction, and blue hues indicate expansion. Purple background indicates non-monotonic behavior. (**B**) Identical response of GS linkers to individual solutes contrasts the differential response of other sequences shown in **Fig. 2B**. Each panel point is the average of the solution-induced change in $\chi$ vs. concentration from two repeats of a specific solute for several different constructs. Vertical lines are the spread of the data.

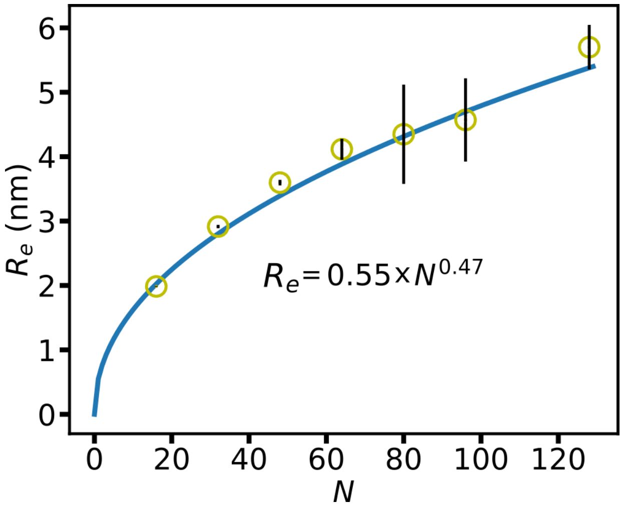


**Figure S2.** End-to-end distance of Gly-Ser repeat sequences as a function of their total number of residues *N*, obtained from all-atom simulations in aqueous solution. Each data point is an average of 5 individual repeats, with lines being the SD of the data. The blue curve is a power law fit of the data, and the power law fit shown in the inset. Our fitted power law, 0.47 ± 0.03 is within error of the power law expected of an ideal polymer (0.5).

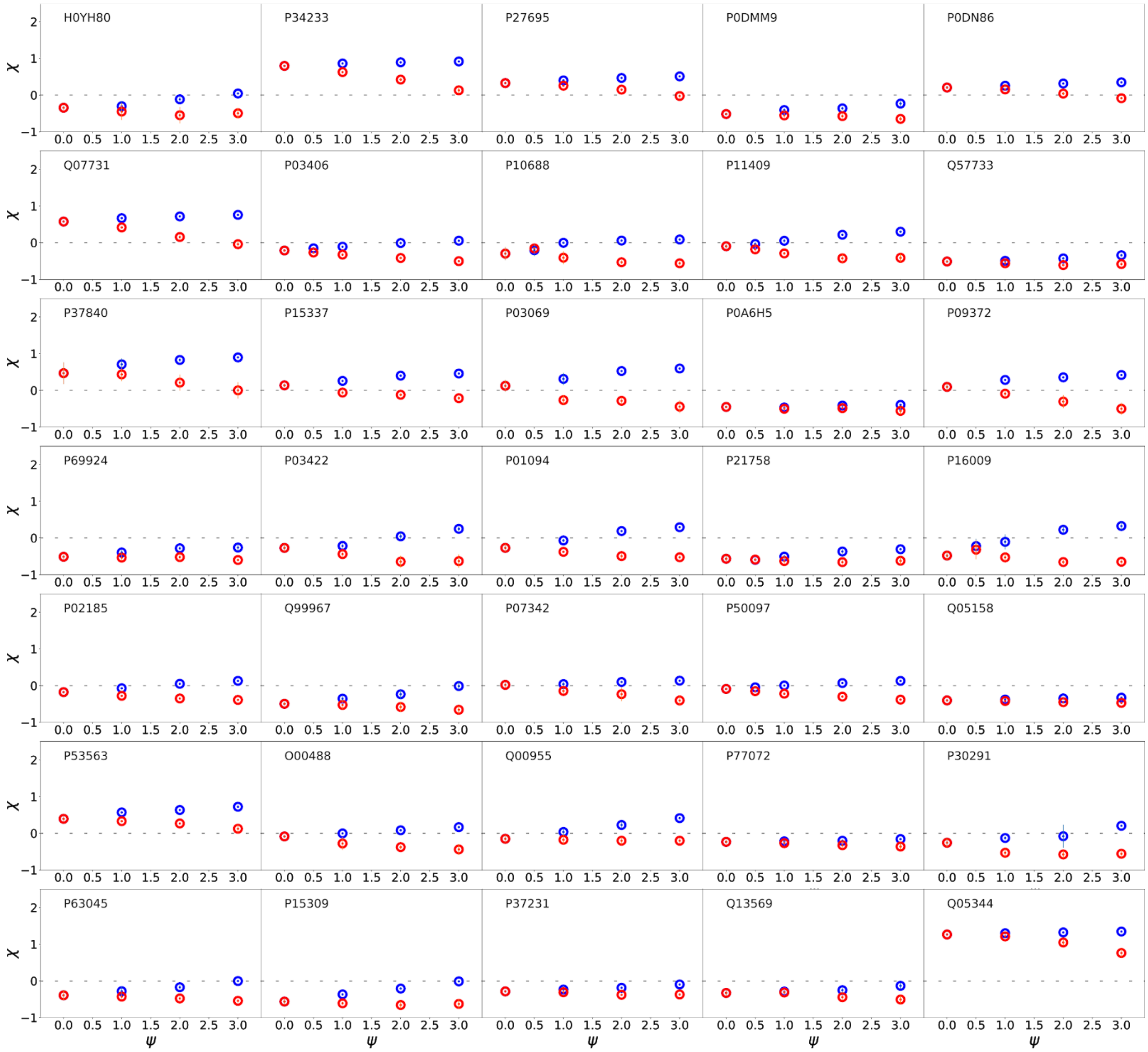


**Figure S3.** $\chi$ vs strength of solution interactions $\psi$ (see **Section S2.1**) for each of the 70 proteins shown in Fig. 3. Each subplot represents a single protein. Blue points are attractive solutions ($\psi > 0$) and red points are repulsive solutions ($\psi < 0$). IDs are UniProt ID when available. All protein names, sequences, and data for each protein are available in **Table S3**.

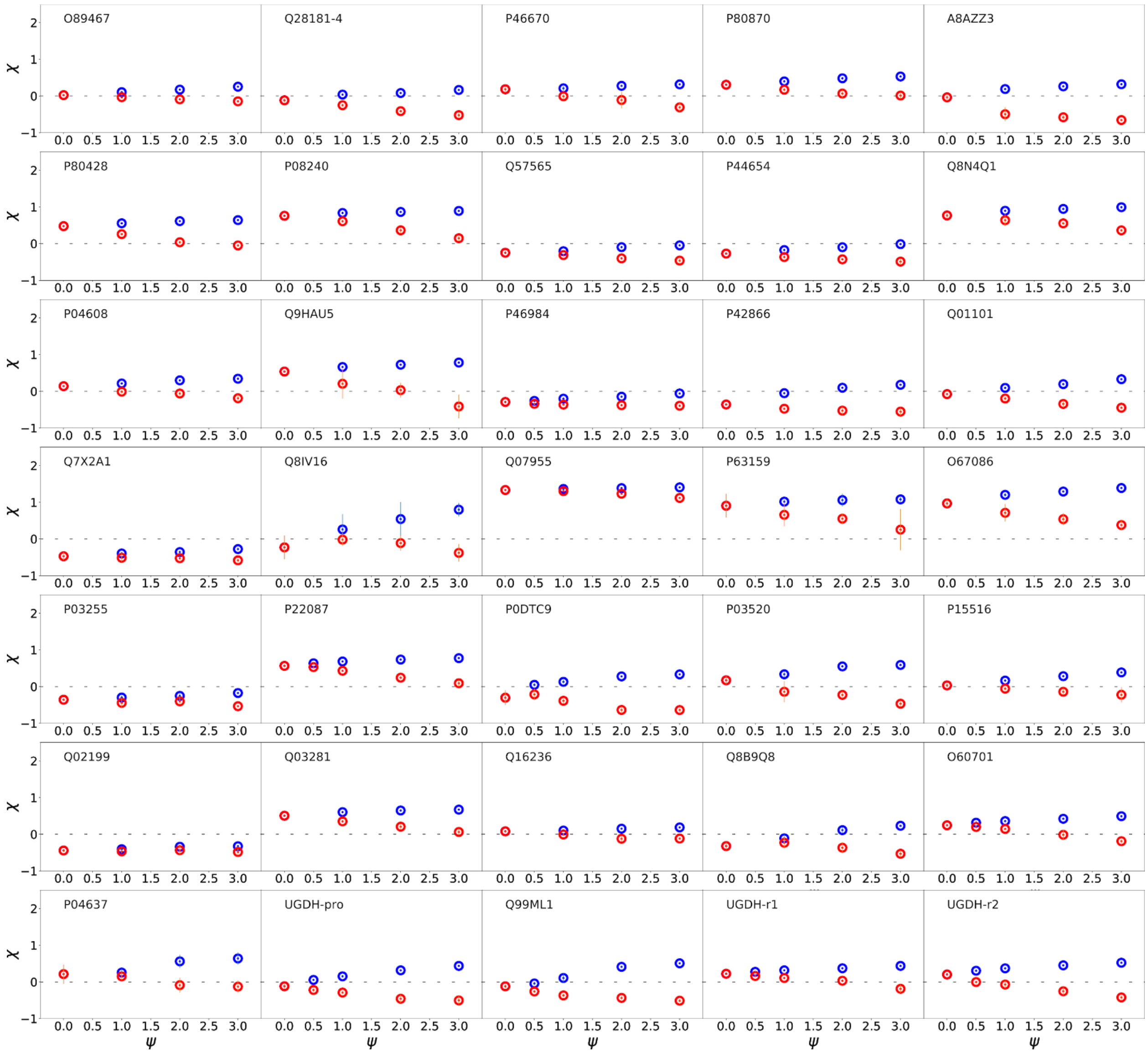


**Figure S3 (cont.).** $\chi$ vs strength of solution interactions $\psi$ (see **Section S2.1**) for each of the 70 proteins shown in Fig. 3. Each subplot represents a single protein. Blue points are attractive solutions ($\psi > 0$) and red points are repulsive solutions ($\psi < 0$). IDs are UniProt ID when available. All protein names, sequences, and data for each protein are available in **Table S3**.

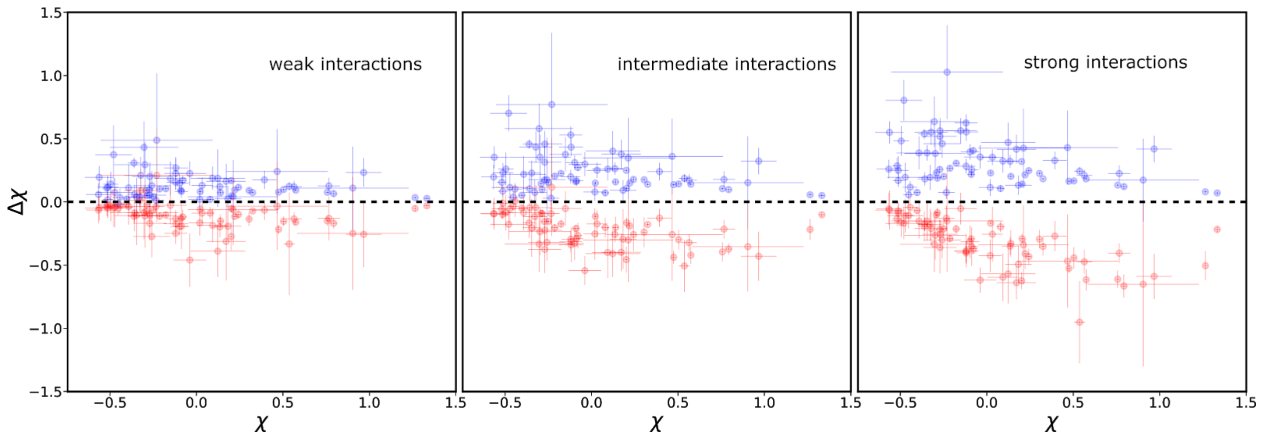


**Figure S4.** Solution sensitivity of proteins shown in **Fig. S3**. Each point represents the solution-induced change in $\chi$ ($\Delta\chi$), for $\psi = \pm1$ (weak interactions), $\pm2$ (intermediate interactions) or $\pm3$ (strong interactions). Blue points represent the response to repulsive solutions and red points represent the response to attractive solutions. Error bars are calculated from 5 independent simulations. See also **Fig. S6**.

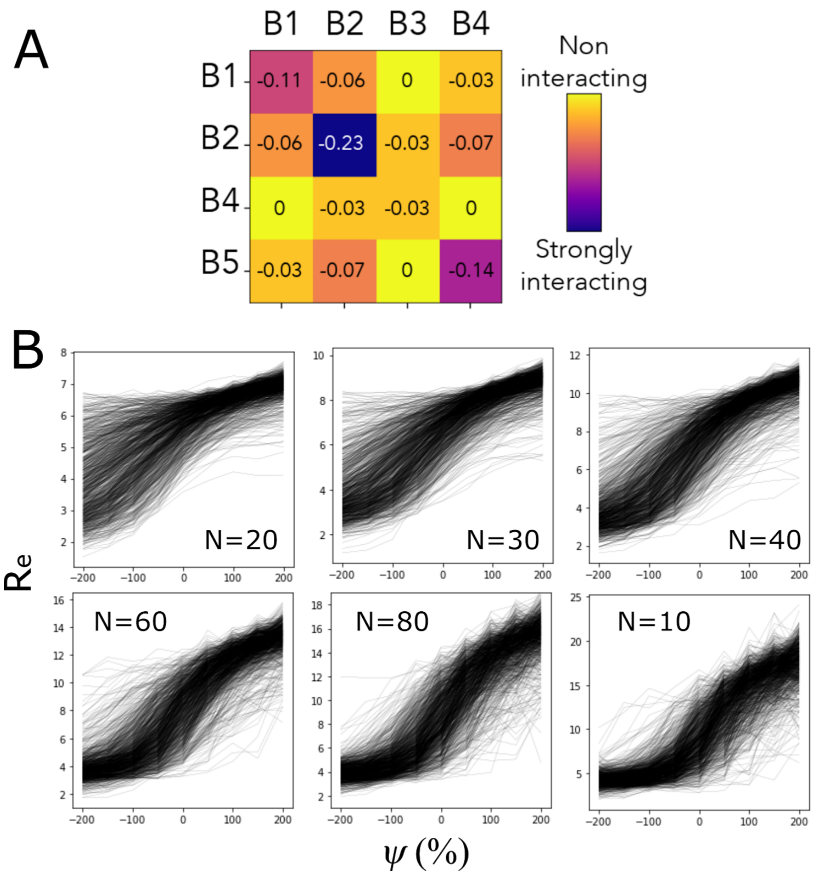


**Figure S5**. **(A)** PIMMS parameters used for heteropolymer simulations. Interaction energies are defined in units of kT and were selected to approximate the chemical diversity observed in polypeptides. B1-B4 are simply "bead" 1 to "bead" 4. **(B)** End-to-end distances (in grid units) for PIMMS coarse-grained simulations of various sequences and chain lengths N. These curves were used to produce the figure shown in **Fig. 4B**.

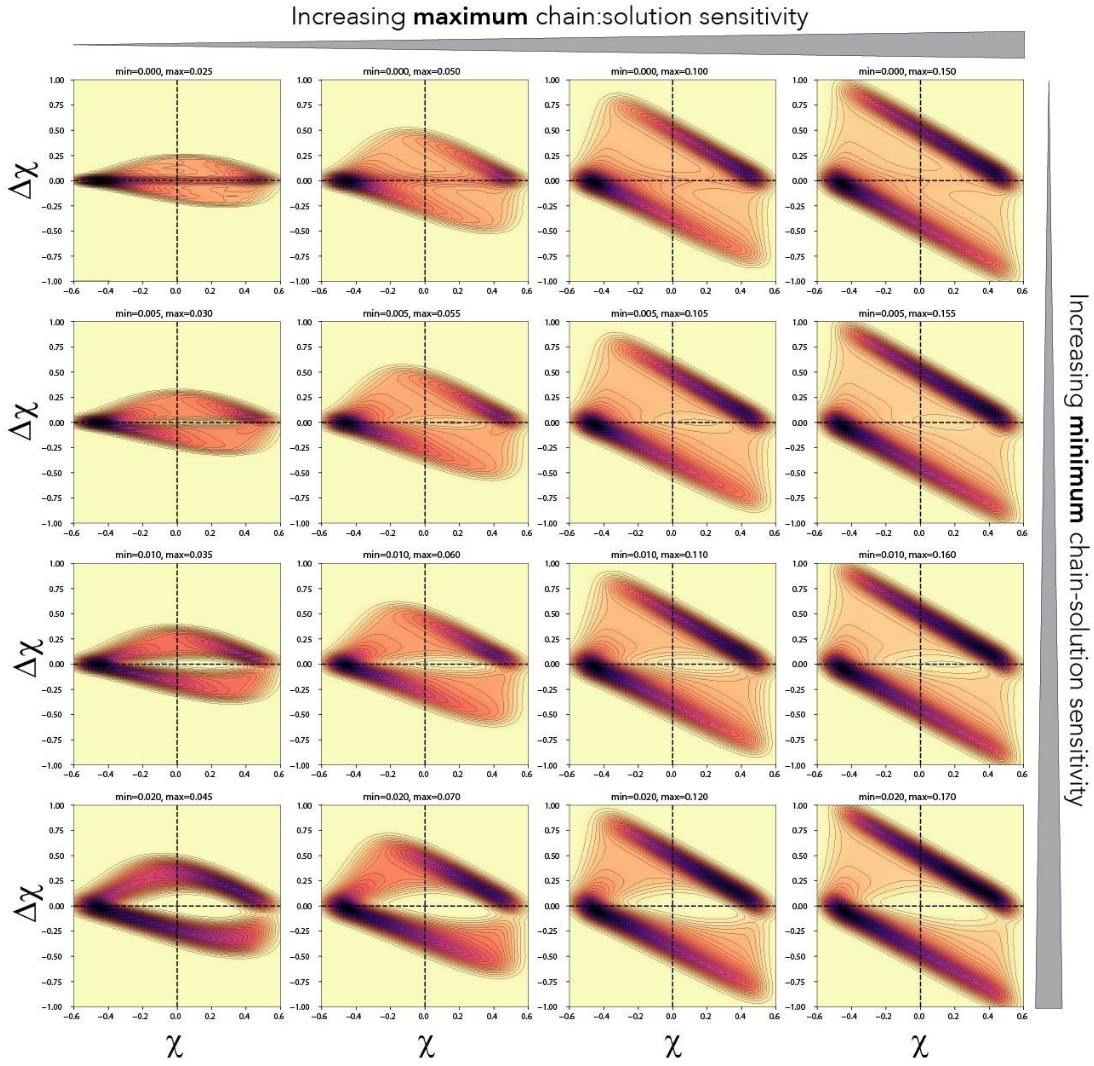


**Figure S6**. Dependence of $\Delta\chi$ vs. $\chi$ as a function of the most and least sensitive chains in an ensemble of sequences. Each figure defines the maximum and minimum perturbation to the chain-solvent interaction. As the maximum perturbation grows (left to right), $\Delta\chi$ becomes larger in a uniform manner along the $\chi$ axis. As the minimum perturbation grows (top to bottom), the opening of a central "pore" region emerges. These two phenomena can be understood intuitively. At the limit of the minimum perturbation being zero, this effectively means there exist chains that are fully insensitive to changes in the solution, such that $\Delta\chi$ is zero. As that minimum increases, *every* chain is somewhat sensitive, with a minimum sensitivity defined by this minimum value. Chains along the coil-to-globule transition are more sensitive than at the coil or globule limits (**Fig. 4D**) such that the pore is centered around $\chi = 0$. The maximum perturbation defines the magnitude of $\Delta\chi$, but is bounded by the chain dimensions, such that $\Delta\chi$ has upper and lower bounds. As the maximum is increased, more perturbations push up against that maximum, such that increasing $\Delta\chi$ density is observed at the bounds (i.e., see top right).

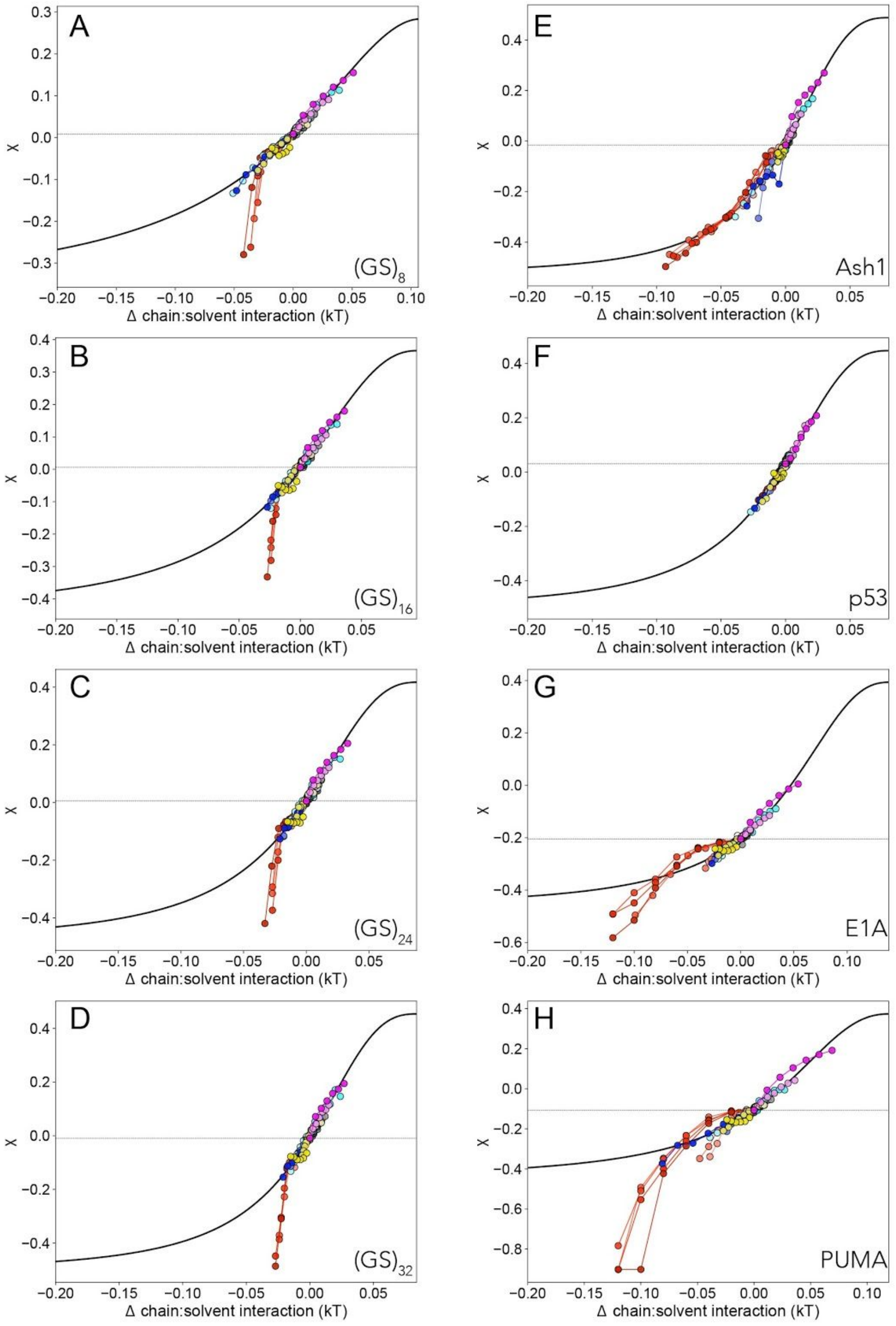


**Figure S7**. Full fit curves for all eight proteins. Horizontal dashed lines reflect the χ value as measured in buffer. Black curve is a length-derived prediction from our analytic model. Note that for many of the curves the high-molecular weight PEG solutions lead to substantial deviations from the master curve, as expected as chain behavior enters the semidilute regime[17], the concentration regime in which PEG chains begin to overlap with one another. PUMA shows the worst agreement with the analytical model, a behavior interpreted as being due to its considerable residual helical structure.

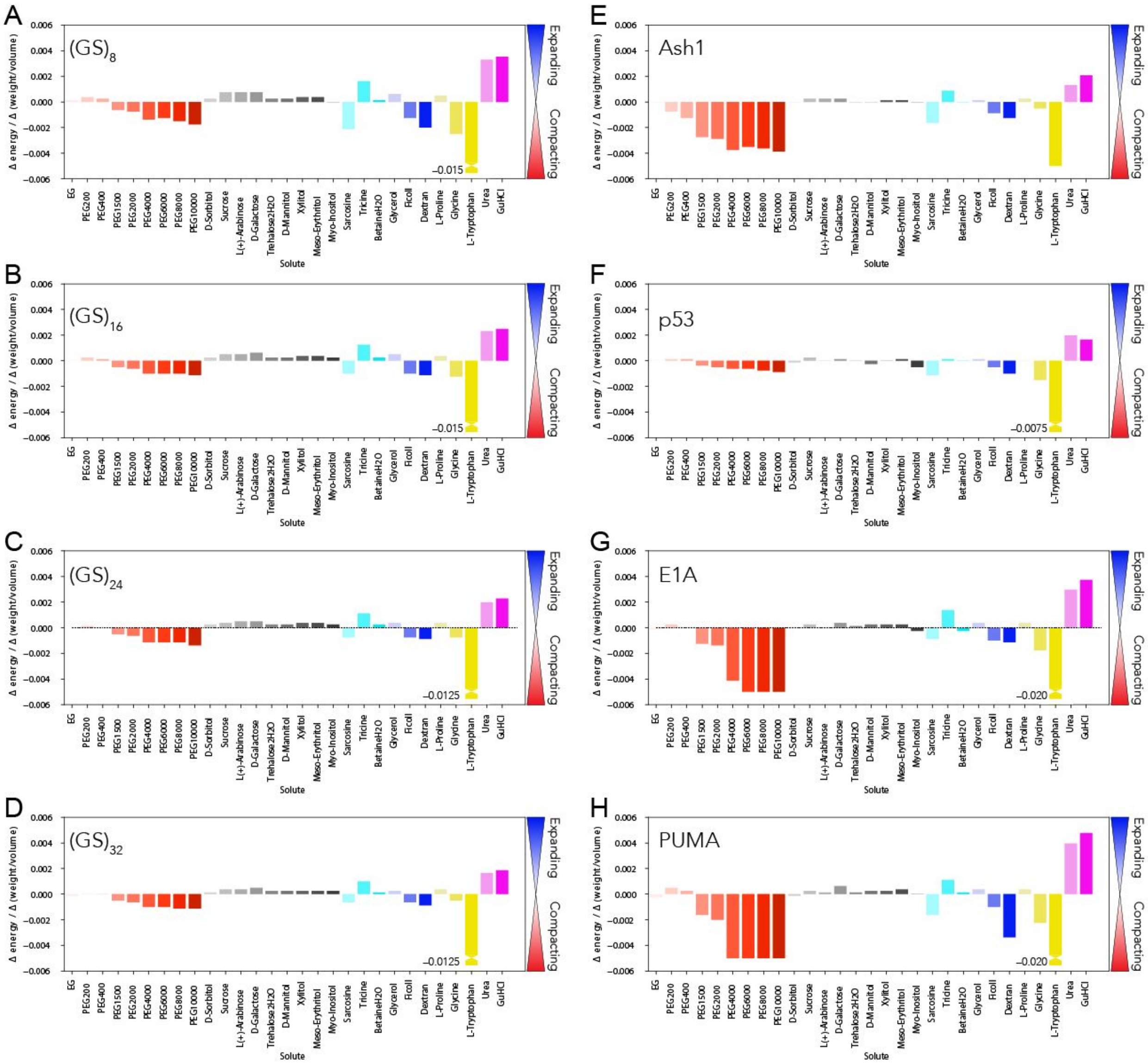


**Figure S8**. Derived solute-specific scalar factors that relate change in chain-solute interaction strength to a change in $\chi$. More positive values lead to chain expansion while more negative values lead to chain compaction.

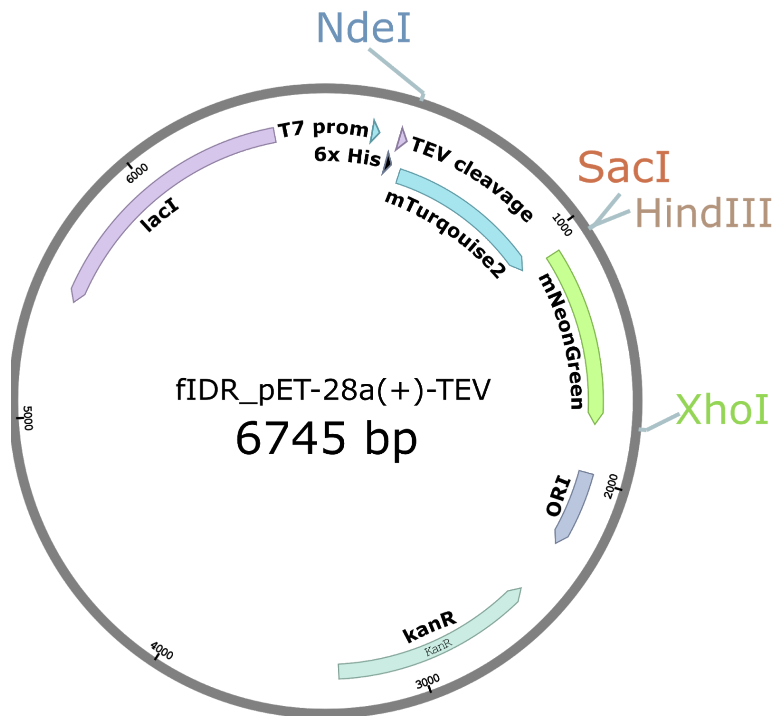


**Figure S9.** Plasmid map for FRET construct bacterial expression vector. Disordered sequences from **Table S4** are inserted between 5' SacI and 3' HindIII restriction sites.

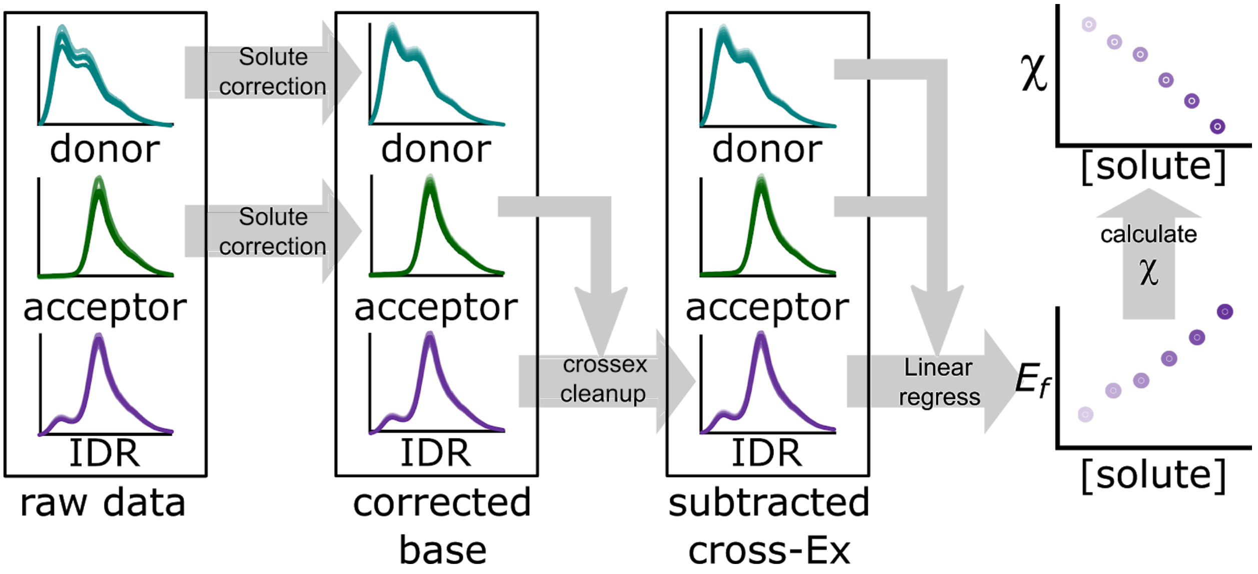


**Figure S10.** Visual summary of the data processing procedure detailed in **SI Section 1.5**. All panels show intensity vs. wavelength data for solutions containing donor-only, acceptor-only, and IDR construct (unless specified otherwise). Spectra are arranged from light to dark going from buffer to high concentrations of solute. Beginning from raw data, base spectra are corrected for pipetting error and protein absorbance to the plate to get corrected base spectra. The acceptor channel is then subtracted from the raw IDR data to remove cross-excitation artifacts. After this, both corrected base spectra are used to fit the corrected IDR spectrum by linear regression. Results of the linear regression are used to calculate the FRET efficiency, $E_f$, as described in **SI Section 1.5**, and $E_f$ is used to calculate $\chi$ as described in **SI Section 1.7**.

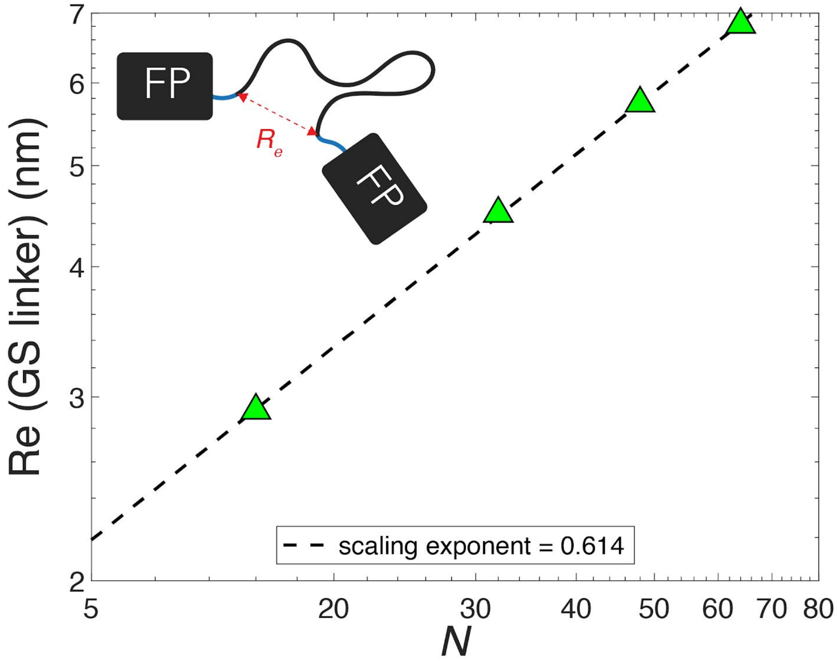


**Figure S11.** Intrachain distance of glycine-serine (GS) linkers connecting two fluorescent proteins in a system that rigorously behaves as a self-avoiding random coil. GS linker end-to-end distance is measured between the first and last residue in the GS repeat region. Note that short (3-7 residue) cloning scars are also present in our model to replicate the actual experimental construct, and these do not contribute residues to the GS linkers in this analysis. Cloning scars are shown as teal parts of the linker.

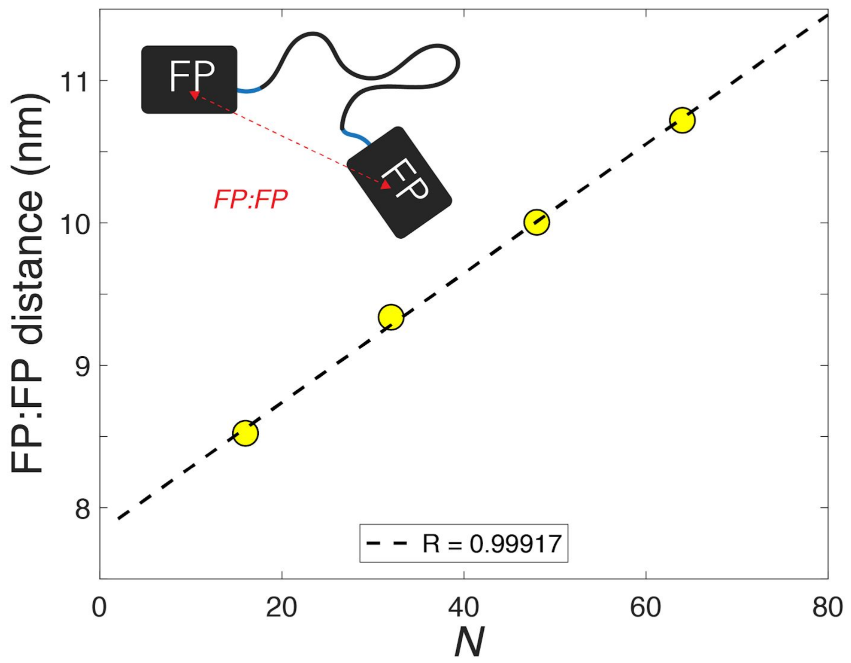


**Figure S12.** Intra-fluorescent protein distance for the same system as in **Fig. S11**. The distance here is measured between the two chromophore centers in each of the two fluorescent proteins. Note that when intra-fluorescent protein distances are measured, we obtain a linear relationship (as opposed to a power law relationship as in **Figs. S11** and **S2**).

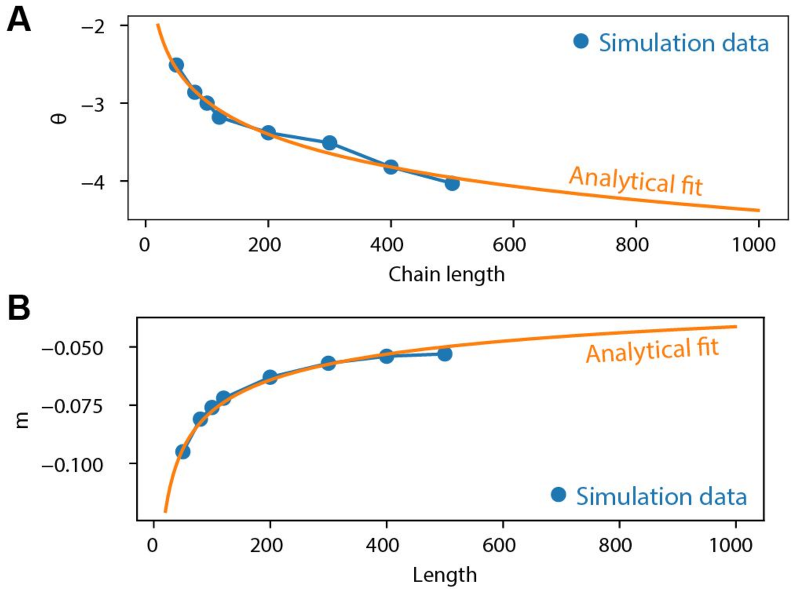


**Figure S13.** Fit of length-dependent model parameters to match PIMMS homopolymer simulations. The orange curves represent the analytical expressions defined in the **Supplementary Methods** using the best fit parameters to fit to the experimentally measured values. **(A)** The fitting of the parameters $c$ and $d$ to reproduce the experimentally-derived length dependence of the cooperativity of the coil-to-globule transition, as quantified by $\theta$. **(B)** The fitting of the parameter $\gamma$ to reproduce the experimentally-derived length dependence of the midpoint on the coil-to-globule transition, as quantified by $m$.

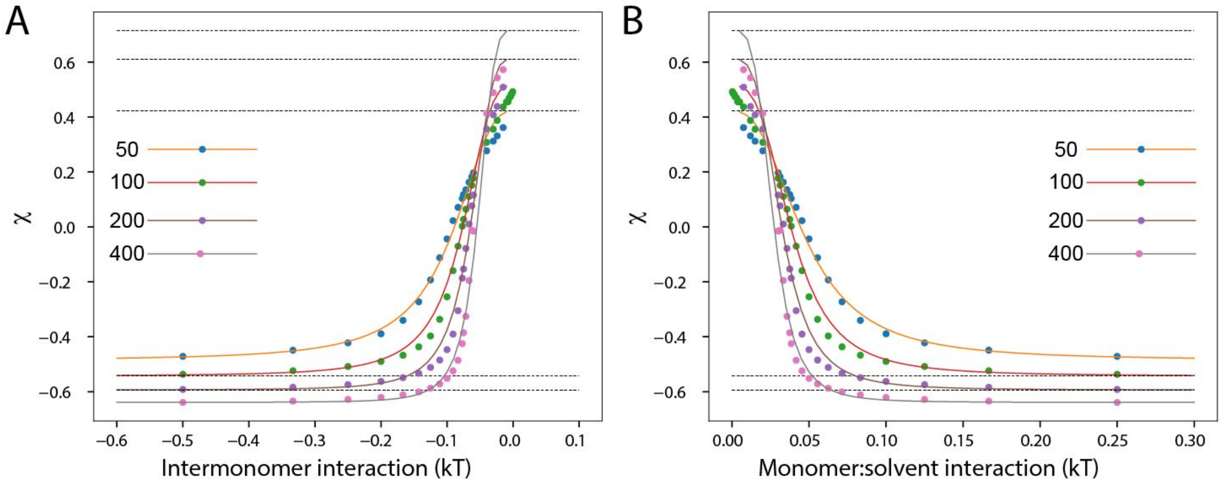


**Figure S14**. Best fit of floating parameters for analytical model (line) to PIMMS simulations (filled circles). (**A)** Data plotted in terms of inter-monomer interaction strength (assuming neutral chain-solvent interactions). **(B)** Same data plotted in terms of chain-solvent interaction strength (assuming neutral inter-monomer interactions). Colors denote chain lengths as specified in the legends.

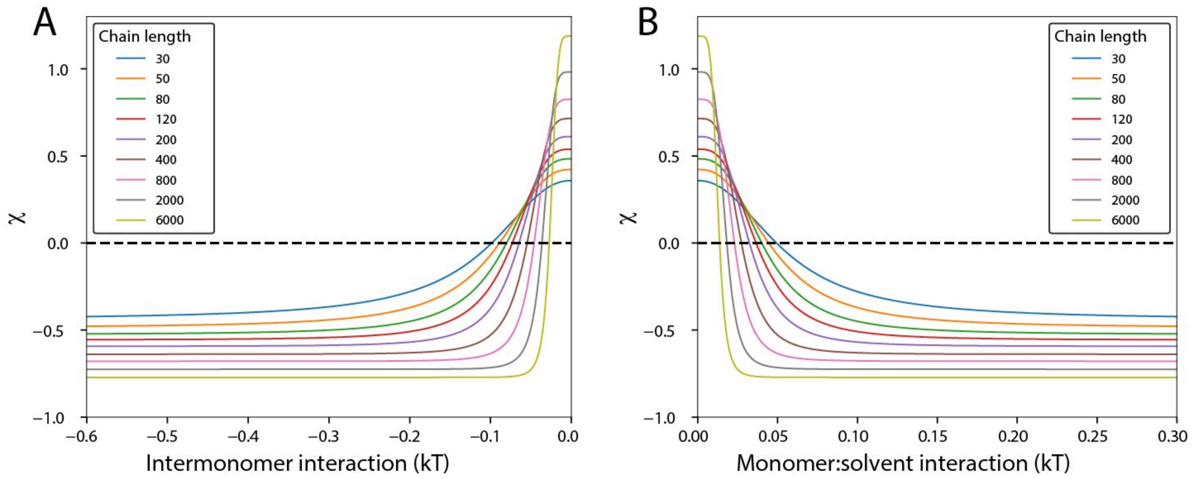


**Figure S15.** Relationship between inter-monomer interaction strength and $\chi$. As chain length increases cooperativity of the coil-to-globule transition increases. Note that the maximum and minimum $\chi$ values show a modest but well-defined length dependence. (**A)** Data plotted in terms of inter-monomer interaction strength (assuming neutral chain-solvent interactions). **(B)** Same data plotted in terms of chain-solvent interaction strength (assuming neutral inter-monomer interactions).

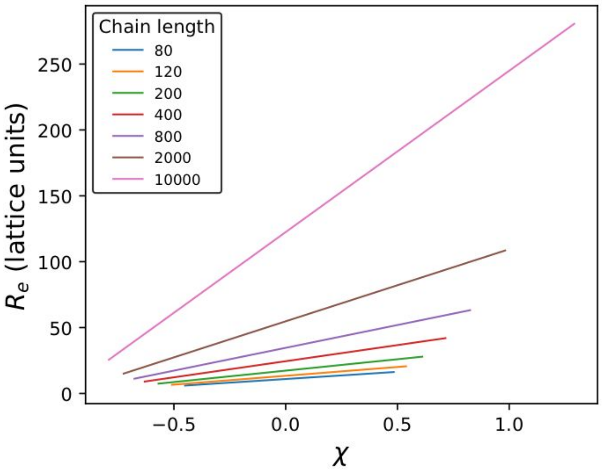


**Figure S16.** Dependence of end-to-end distance ($R_e$) on $\chi$. As the chain becomes longer both the maximum and the steepness of the $R_e$-dependence on $\chi$ becomes larger.

### SI References


(1) Cranfill, P. J.; Sell, B. R.; Baird, M. A.; Allen, J. R.; Lavagnino, Z.; de Gruiter, H. M.; Kremers, G.-J.; Davidson, M. W.; Ustione, A.; Piston, D. W. Quantitative Assessment of Fluorescent Proteins. *Nat. Methods* **2016**, *13* (7), 557–562.
(2) Lambert, T. J. FPbase: A Community-Editable Fluorescent Protein Database. *Nat. Methods* **2019**, *16* (4), 277–278.
(3) Mastop, M.; Bindels, D. S.; Shaner, N. C.; Postma, M.; Gadella, T. W. J., Jr; Goedhart, J. Characterization of a Spectrally Diverse Set of Fluorescent Proteins as FRET Acceptors for mTurquoise2. *Sci. Rep.* **2017**, *7* (1), 11999.
(4) Rubinstein, M.; Colby, R. H. *Polymer Physics*; Oxford University Press: New York, 2003.
(5) Peran, I.; Holehouse, A. S.; Carrico, I. S.; Pappu, R. V.; Bilsel, O.; Raleigh, D. P. Unfolded States under Folding Conditions Accommodate Sequence-Specific Conformational Preferences with Random Coil-like Dimensions. *Proc. Natl. Acad. Sci. U. S. A.* **2019**, *116* (25), 12301–12310.
(6) Holehouse, A. S.; Garai, K.; Lyle, N.; Vitalis, A.; Pappu, R. V. Quantitative Assessments of the Distinct Contributions of Polypeptide Backbone Amides versus Side Chain Groups to Chain Expansion via Chemical Denaturation. *J. Am. Chem. Soc.* **2015**, *137* (8), 2984–2995.
(7) Mohsen-Nia, M.; Modarress, H.; Rasa, H. Measurement and Modeling of Density, Kinematic Viscosity, and Refractive Index for Poly (ethylene Glycol) Aqueous Solution at Different Temperatures. *J. Chem. Eng. Data* **2005**, *50* (5), 1662–1666.
(8) Vitalis, A.; Pappu, R. V. ABSINTH: A New Continuum Solvation Model for Simulations of Polypeptides in Aqueous Solutions. *J. Comput. Chem.* **2009**, *30* (5), 673–699.
(9) Mittal, A.; Das, R.; Vitalis, A.; Pappu, R. ABSINTH Implicit Solvation Model and Force Field Paradigm for Use in Simulations of Intrinsically Disordered Proteins. *Computational Approaches to Protein Dynamics: From Quantum to Coarse-Grained Methods* **2014**, 181.
(10) Holehouse, A. S.; Sukenik, S. Controlling Structural Bias in Intrinsically Disordered Proteins Using Solution Space Scanning. *J. Chem. Theory Comput.* **2020**. https://doi.org/10.1021/acs.jctc.9b00604.
(11) Piovesan, D.; Tabaro, F.; Mičetić, I.; Necci, M.; Quaglia, F.; Oldfield, C. J.; Aspromonte, M. C.; Davey, N. E.; Davidović, R.; Dosztányi, Z.; Elofsson, A.; Gasparini, A.; Hatos, A.; Kajava, A. V.; Kalmar, L.; Leonardi, E.; Lazar, T.; Macedo-Ribeiro, S.; Macossay-Castillo, M.; Meszaros, A.; Minervini, G.; Murvai, N.; Pujols, J.; Roche, D. B.; Salladini, E.; Schad, E.; Schramm, A.; Szabo, B.; Tantos, A.; Tonello, F.; Tsirigos, K. D.; Veljković, N.; Ventura, S.; Vranken, W.; Warholm, P.; Uversky, V. N.; Dunker, A. K.; Longhi, S.; Tompa, P.; Tosatto, S. C. E. DisProt 7.0: A Major Update of the Database of Disordered Proteins. *Nucleic Acids Res.* **2017**, *45* (D1), D219–D227.
(12) McGibbon, R. T.; Beauchamp, K. A.; Harrigan, M. P.; Klein, C.; Swails, J. M.; Hernández, C. X.; Schwantes, C. R.; Wang, L.-P.; Lane, T. J.; Pande, V. S. MDTraj: A Modern Open Library for the Analysis of Molecular Dynamics Trajectories. *Biophys. J.* **2015**, *109* (8), 1528–1532.
(13) Martin, E. W.; Holehouse, A. S.; Peran, I.; Farag, M.; Incicco, J. J.; Bremer, A.; Grace, C. R.; Soranno, A.; Pappu, R. V.; Mittag, T. Valence and Patterning of Aromatic Residues Determine the Phase Behavior of Prion-like Domains. *Science* **2020**, *367* (6478), 694–699.
(14) Hofmann, H.; Soranno, A.; Borgia, A.; Gast, K.; Nettels, D.; Schuler, B. Polymer Scaling Laws of Unfolded and Intrinsically Disordered Proteins Quantified with Single-Molecule

Spectroscopy. *Proceedings of the National Academy of Sciences* **2012**, *109* (40), 16155–16160.

(15) Holehouse, A. S.; Pappu, R. V. Collapse Transitions of Proteins and the Interplay Among Backbone, Sidechain, and Solvent Interactions. *Annu. Rev. Biophys.* **2018**, *47*, 19–39.

(16) Sørensen, C. S.; Kjaergaard, M. Effective Concentrations Enforced by Intrinsically Disordered Linkers Are Governed by Polymer Physics. *Proc. Natl. Acad. Sci. U. S. A.* **2019**, *116* (46), 23124–23131.

(17) Kozer, N.; Kuttner, Y. Y.; Haran, G.; Schreiber, G. Protein-Protein Association in Polymer Solutions : From Dilute to Semidilute to Concentrated. *Biophys. J.* **2007**, *92* (6), 2139–2149.
